# Prediction of harvest-related traits in barley using high-throughput phenotyping data and machine learning

**DOI:** 10.1101/2025.05.29.656856

**Authors:** Hans Tietze, Lamis Abdelhakim, Barbora Pleskačová, Ayelet Kurtz-Sohn, Eyal Fridman, Zoran Nikoloski, Klára Panzarová

## Abstract

Developing crop varieties that maintain productivity under drought is essential for future food security. Here, we investigated the potential of time-resolved high-throughput phenotyping to predict harvest-related traits and identify drought-stressed plants. Six barley lines (*Hordeum vulgare*) were grown in a greenhouse environment with well-watered and drought treatments, and phenotyped using RGB, thermal infrared, chlorophyll fluorescence and hyperspectral imaging sensors. Temporal phenomic classification model accurately distinguished between drought-treated and control plants, achieving high accuracy (R^2^ ≥ 0.97) even when exclusively using predictors only from the early phase after drought induction. Canopy temperature depression at the early stage and RGB-derived plant size estimates at the late stage were identified as key classification features. Temporal phenomic prediction model of harvest-related traits achieved particularly high mean R^2^ values for total biomass dry weight (0.97) and total spike weight (0.93), with RGB plant size estimators emerging as important predictors. Prediction accuracy for these traits remained high (R^2^ ≥ 0.84) when using only predictors from the first half of the experiment. Models trained on pooled drought and control data outperformed single-treatment models and retained high accuracy when applied across treatments. These findings support the integration of high-throughput phenotyping and temporal modelling to enable timely and more cost-effective selection of drought-resilient genotypes, and illustrate the broader potential of phenomics-driven approaches in accelerating crop improvement under stress-prone conditions.

## Introduction

Climate change influences agricultural productivity and negatively affects crop yield, making the breeding of resilient crop varieties essential. The development of such stress-resilient varieties is challenging due to the interaction between genotype and environment that shapes complex traits, like grain yield. As a result, enhancing breeding programs for resilient crops requires accurate yield prediction across diverse environments (Cooper & Messina, 2023). Developing predictive models that integrate diverse data sets, e.g., genomics along with spatiotemporal phenomics and enviromics, can support this goal by enabling more accurate prediction of crop phenotypes (Xu et al., 2022). In addition, the implementation of advanced breeding techniques demands the development and deployment of high-throughput phenotyping (HTP) platforms. The resulting data along with the computational and machine learning approaches can improve future yield performance and help in developing resilient crop varieties that can withstand a variety of stresses, typical of field conditions (Varshney et al., 2021).

HTP is one of the techniques that has transformed and accelerated plant breeding by enabling large-scale, rapid screening of different phenotypic traits of interest, including automated data acquisition and trait analysis (Song et al., 2021; Yang et al., 2020). The use of multi-imaging sensors is essential for the non-invasive and precise assessment of plant growth dynamics as well as physiological responses. This approach provides a comprehensive view of plant development, enabling the monitoring and assessment of plant performance and stress responses (Humplík et al., 2015). Many studies have investigated the effect of abiotic stress, including drought, aiming at identifying the key phenotypic traits and physiological mechanisms that enhance stress tolerance (Al-Tamimi et al., 2022; Findurová et al., 2023). However, the complex nature of genotype-by-environment interactions remains a major challenge and demands further investigation. Moreover, a better understanding of drought adaptation requires recognizing that the impact of stress on physiological traits linked to grain yield can vary depending on stress intensity, genotype susceptibility, and developmental stage (Khadka et al., 2020). Along these lines, advances in high-throughput and precision phenotyping techniques have contributed to improve the strategies for mitigating the adverse effects of drought stress on plants and enhancing their resilience and productivity (Farooq et al., 2024).

One of the main challenges in harnessing the potential of high-throughput data lies in the management and analysis to identify traits of interest and reveal plant responses to stress (Langstroff et al., 2022; Leonelli et al., 2017). Data generated at multiple spatial and temporal scales requires robust analytical pipelines capable of handling such complex phenotypic datasets (Tardieu et al., 2017). Moreover, in phenotyping studies focusing on stress response across developmental stages, models are often modified to capture the dynamic changes of plant response to stress over time (Li et al., 2020). Recent pioneering advances have facilitated the prediction of the dynamics of multiple traits given genetic markers alone (Hobby et al., 2025).

Machine learning techniques play a transformative role in phenotypic data analysis by linking large, complex datasets to traits of interest (Singh et al., 2016). Combining image-based phenotyping with machine learning approaches has enabled the extraction of new insights from curated, annotated, high-dimensional data sets across various crops and stress conditions (Singh et al., 2021). Machine learning encompasses a range of techniques, including feature extraction, pattern recognition, classification, and prediction. Some of these approaches facilitate the analysis of complex phenotypic data sets by considering multiple traits simultaneously, accounting for trait integration (Mbebi et al., 2025). As such, applying machine learning to phenomic data provides a powerful framework for uncovering patterns and extracting biologically meaningful insights (Gill et al., 2022).

In this study, we used an HTP platform equipped with multiple imaging sensors. We aimed to develop an advanced data analysis pipeline and apply it to perform a phenotypic data analysis of different barley (*Hordeum vulgare*) lines exposed to drought stress. We focused on barley as it is a model cereal crop (FAO, 2023; Newton et al., 2011), and we aimed to investigate the impact of drought as a predominant stress in future climate scenarios (IPCC, 2021). This was achieved by: (i) using a classification model to identify distinct traits that differentiate drought-stressed from well-watered plants and (ii) using regression models to accurately predict harvest-related traits. The applied modelling approach enabled pinpointing the most predictive traits at specific time points. Moreover, early detection of such traits can support breeders in selecting stress-tolerant genotypes more efficiently, potentially accelerating the development of resilient crop varieties and improving resource use in breeding programs.

## Materials and methods

### Plant material and growth conditions

Six genetically homogenous barley lines were selected in this study, including one elite cultivar line (Barke) (L1) and five lines originating from the CMPP (Cytonuclear MultiParent Population) (L2-L5) and HEB-25 (Halle exotic barley) (L6) populations (HÜbner et al., 2009) (Supp. Table S1).

After seeds were stratified at 4°C in darkness, seedlings were transferred to light in the walk-in chamber (FytoScope FS-WI, Photon Systems Instruments (PSI), Drásov, Czech Republic) and were grown under a short-day regime, until the emergence of the fifth leaf. One seedling was transplanted per 3L pot filled with 1850 g of Klasmann Substrate-2 : sand (3:1). Plants were transferred to the greenhouse under a long day regime (16h photoperiod), 22 ± 3 / 17 ± 2 °C for day/night temperature, and 51 ± 8 / 62 ± 4 % for day/night relative humidity.

### Phenotyping protocol

The experiment was conducted in a greenhouse that is connected to the PlantScreen^TM^ Modular phenotyping platform (PSI, Czech Republic), where pots were placed on transportation disks carried from the growth area toward the multi-imaging and irrigation units. Plant performance, including morphological and physiological responses, was assessed throughout the whole life cycle with an overall duration of plant cultivation of 97 days after transfer to light (DAT), and kept until reaching the full maturation stage (126 DAT). Over the course of 10 weeks, the daily phenotyping protocol was conducted to extract morpho-physiological and spectral-related traits in plants cultivated in semi-controlled greenhouse conditions under two watering regimes, control and progressive drought stress regime. Drought-stressed plants were maintained at 25% soil relative water content (SRWC) till the flowering stage, then watering was further reduced to 20% SRWC. We used nine biological replicates per treatment for most of the lines, and 20 replicates per treatment for the HEB line and elite line (Barke), which served as the reference line. The reduced watering regime was induced at the tillering stage (24 DAT) and remained reduced for the stressed group for the whole cultivation period. On a daily basis, plants were randomized in the cultivation greenhouse to avoid positional effects, environmental conditions were recorded with minute resolution, and daily watering and weighing of the plants were performed. Plants were phenotyped daily up to the maturity stage using multi-imaging sensors of PlantScreen^TM^ Modular phenotyping platform (PSI, Czech Republic), including RGB, thermal infrared (IR), chlorophyll fluorescence and hyperspectral imaging, as described in (Abdelhakim et al., 2024). Referring to chlorophyll fluorescence imaging, different measuring protocols were selected for capturing more insights into the photosynthetic performance, including a morning protocol and two different evening protocols. During the day (light-adapted state), measuring protocols were optimized to measure the quantum yield of PSII (QY_Lss) under two light steady state (Lss) intensities, including high light (HL, Lss1 at 1200 µmol.m^-2^s^-1^) and low light (LL, Lss2 at 130 µmol.m^-2^s^-1^). To estimate the plasticity index of QY under different light intensities, the ratio between QY_Lss measured under low (Lss2) to high (Lss1) light was calculated. Moreover, measurement on dark-adapted plants was conducted to assess the photosynthesis induction and relaxation kinetics during the night period at two different light intensities protocols, i.e., High light (HL) at 1200 µmol.m^-2^s^-1^ and conditional light (CL) at 360 µmol.m^-2^s^-1^ (Fig. 1). At the end of the maturation stage, the total biomass of the plants was manually harvested, including analysis of the total tiller number, spike number, and other spike-related traits (Supp. Table S2).

**Figure 1.**
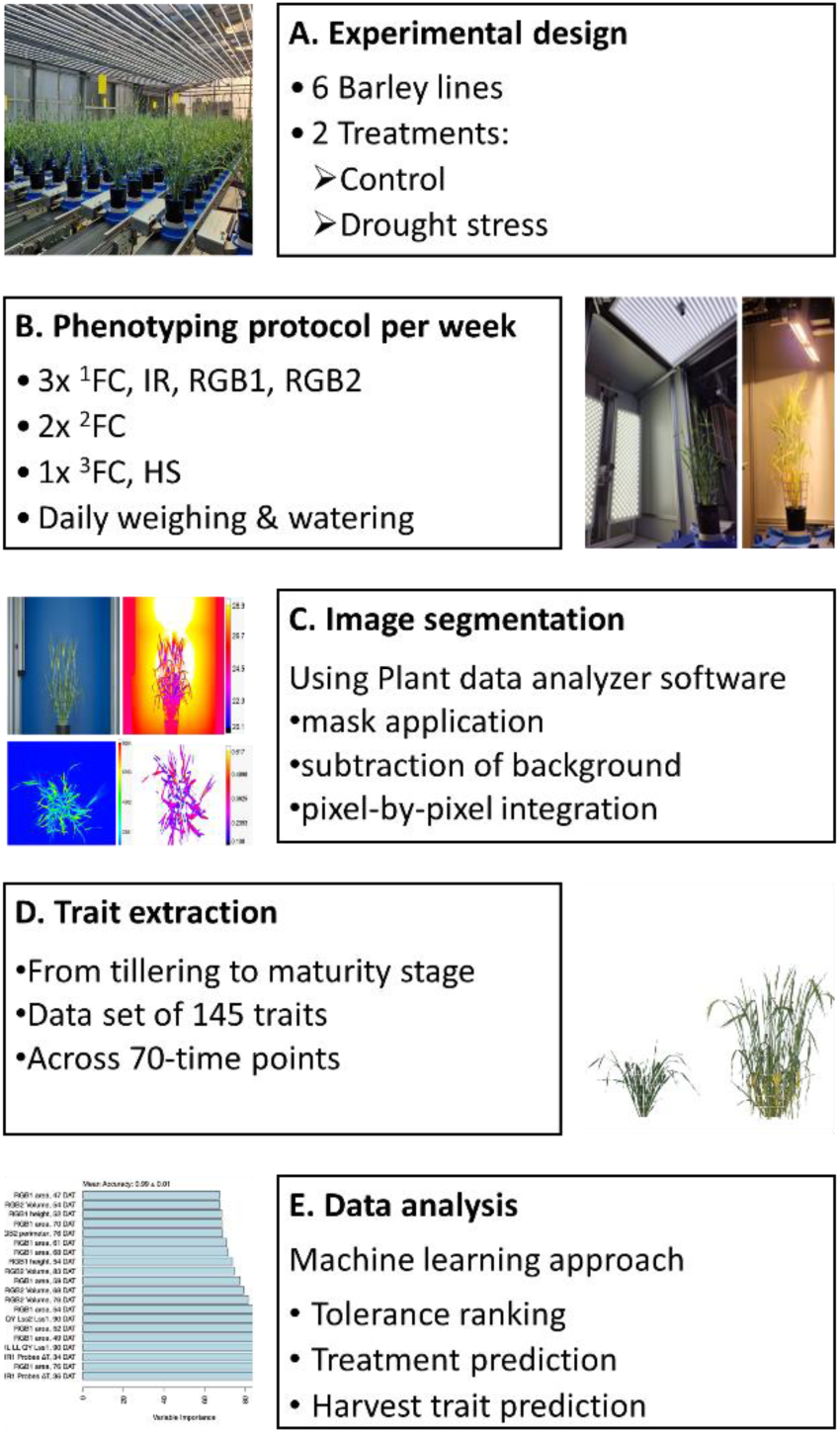
Overview of the experimental design and measurements performed. A. Summary of the experimental design, including 6 barley lines. Two different water regime treatments were applied, control and drought stress at 60 % and 25-20% soil relative water content, respectively. Phenotyping was conducted from the tillering stage to the maturity stage, followed by final harvest. B) Automated image-based phenotyping using the PlantScreen^TM^ Modular phenotyping platform at PSI Research Center, where plants are moved from the greenhouse growing area towards imaging units. The phenotyping protocol was conducted daily with different protocols. In chlorophyll fluorescence imaging using FluorCam (FC), ^1^FC morning measurement, ^2^FC night measurement, ^3^FC chlorophyll content were conducted, thermal infra-red imaging (IR), RGB^3^ including two angles from RGB side and one angle RGB top view, and hyperspectral imaging (HS) including SWIR and VNIR imaging. C. Automated image segmentation process. D. Extracting traits, including measured and calculated parameters, among the developmental stages. E. Data analysis using a machine learning approach to assess tolerance of plants under drought stress, discriminate between the two water regimes and find the most predictive traits of the final yield.

### Data processing pipeline

The gathered dataset consists of dynamic phenotypical data from 70 time points captured for the six barley lines that were grown under two conditions, including 9-20 replicates per treatment per line. Overall, 145 image-based and post-harvest traits were extracted (Supp. Table S2) and subjected to further data analysis. Of these, 52 traits from chlorophyll fluorescence and thermal IR imaging were excluded from downstream analysis, as they represented raw measurements used solely in the calculation of more biologically meaningful derived indices. The full data analysis pipeline was conducted using R studio (version 4.3.2).

Due to differing assumptions about input data across methods, pre-processing followed multiple branching paths. For analyses that separated drought and control treatments (harvest prediction per treatment), data were partitioned before outlier detection and transformation. In contrast, pooled-treatment analyses (i.e., variance decomposition, treatment classification, pooled-treatment harvest prediction) preserved treatment-induced variance by avoiding such partitioning. For temporal traits, each measurement time point was treated as a separate data group.

To maximize sample size and model robustness for the genotype-agnostic methods, including temporal phenomic classification (TPC) and prediction (TPP), three additional genotypic lines (L7-L9) were used in part of the analysis pipeline with the six genetically homogeneous lines (L1-L6) (Supp. Table S1). However, as a result of heterogeneity, those three lines were excluded from analyses that assumed genetic homogeneity (i.e., clustering of samples, drought tolerance ranking, and trait variance decomposition).

### Handling of missing values

Data pre-processing began with the imputation of 10 missing values across 267 plant samples in harvest-traits. Missing values were found in spike weight (nine samples) and total biomass dry weight (one sample). These were evenly distributed among samples, except for one sample (L8_D_15, Drought) with missing values in two traits. Missing values were imputed using MissForest imputation (Stekhoven & Bühlmann, 2011), implemented using the R package missForest (version 1.4), leveraging the remaining harvest traits. The out-of-bag (OOB) error for these imputations is provided in Supp. Table S3.

### Outlier handling

Outliers were identified as data points exceeding three times the interquartile range (IQR) of a given data group. These values were removed and re-imputed using missing forest imputation. The OOB errors for these imputations were reported (Supp. Table S4). At this stage, the processed data was exported for variance decomposition.

### Data transformation

A Shapiro-Wilk Test for normality was applied to every data group and p-values were corrected using the Bonferroni method. Those groups whose distribution was deemed non-normal had a Box-Cox transformation and were tested for normality again. Cases of non-normality before and after correction were reported (Supp. Table S5). Following Box-Cox transformation of some groups, a Z-score transformation was applied to all groups. At this point, the transformed data were used for treatment classification and harvest trait prediction.

### Week-wise aggregation of predictors

The same pre-processing steps used with the non-aggregated data set were also employed with the aggregated data set, including: group-wise random forest imputation using missForest, outlier detection based on the IQR with a threshold of 3, re-imputation of extreme values after their removal, and transformation of non-normal trait distributions using Box-Cox followed by z-score normalization. Branching preprocessing paths were also applied, where treatment-specific analyses were conducted on partitioned data, while pooled-treatment analyses preserved treatment-induced variance by processing all samples jointly. A key difference lies in the temporal structuring of the data, whilst the original pipeline treated each measurement daily time point (DAT) as a separate data group, this pipeline uses weekly phases (WP) for grouping and aggregation. This approach reduces temporal noise while maintaining biological resolution, particularly relevant for trait dynamics across stress and recovery phases. The mapping between the DAT and the corresponding WP was defined in Supp. Table S5. For each numeric variable, the average (mean), minimum, and maximum values were calculated. To reduce redundancy, if all three values were identical within a group, indicating no variation, the minimum and maximum columns were removed, leaving only the average as the sole predictor.

### Clustering of samples

All temporal traits from weekly aggregated data were combined, and principal component analysis (PCA) was performed using two R packages, prcomp and PCAtools, with scaled and centered data to explore the underlying structure. Unsupervised clustering using k-means was then applied to the scaled trait data. Trait means were computed for each genotype-treatment combination and scaled. The optimal number of clusters was determined using the silhouette method with the R package factoextra (fviz_nbclust function). Clustering results were visualized using the fviz_cluster function and projected onto PCA space. Clusters were annotated by genotype and treatment, with PCA coordinates overlaid with confidence hulls to enhance interpretability.

### Drought tolerance ranking of lines

The barley lines were ranked based on the magnitude of drought-induced effects on phenotypic traits. A permutational multivariate analysis of variance (PERMANOVA) with 3000 permutations was applied separately to temporal and harvest traits, comparing drought-treated and control plants within each genotype (Anderson, 2017). This analysis was conducted using the R package vegan (version 2.6-8). Generalized eta squared (η²) was used to quantify the treatment effect size, providing a measure of how strongly drought influenced each genotype’s temporal or harvest traits.

### Temporal phenomic classification of treatment

Random Forest (RF) binary classification of plant treatment was performed using temporal traits as predictors. Models were trained using the R packages caret (version 6.0-94) and randomForest (version 4.7-1.1). Each trait at each time point was used as an independent predictor. Models were trained using a 3-fold 5-repeat cross validation (CV) scheme. In each repeat, data points were split randomly into three folds. For each split, two folds were used for training the model, and the last was reserved for testing. This was repeated three times, leaving each fold out for testing once.

To optimize model performance, the number of predictors randomly selected at each decision tree split was treated as a tunable parameter. A range of candidate values was systematically evaluated, and model performance was assessed across the 15 different test sets (3 folds × 5 repeats) to ensure robustness. The final selection was based on the average performance across all repetitions, with the best-performing setting chosen to balance accuracy and model complexity.

Temporal phenomic classification (TPC) of treatment was conducted using temporal traits from all time points (n_predictors_ = 850) and using predictors from each separate week of the experiment (53 ≤ n_predictors_ ≤ 92, depending on the week). Model accuracies were compared using one-way ANOVA and a pairwise t-test with Holm’s method for p-value correction.

To assess the variable importance of predictors, a permutation-based approach was used. For each tree in the forest, the classification accuracy was first recorded using the out-of-bag (OOB) data, which consists of observations left out during bootstrap sampling. Then, the values of a given predictor were randomly permuted in the OOB data, and classification accuracy was re-evaluated. The drop in accuracy due to this permutation, relative to the original OOB accuracy, was computed for each tree. This accuracy difference was averaged across all trees, normalized by the standard deviation of the differences, and then scaled so that the most important variable received an importance score of 100.

### Temporal phenomic prediction of harvest traits

Temporal Phenomic Prediction (TPP) of harvest traits was performed using least absolute shrinkage and selection operator (LASSO) with R package glmnet (version 4.1-8) and RF with R package randomForest (version 4.7-1.1) regression models. Training was done using the R package caret (version 6.0-94). The internal CV schedule was largely equivalent to the one used in TPC, except that LASSO models were optimized for the regularization strength parameter, rather than the number of variables considered at each split. In addition, the 3-fold CV procedure was repeated 15 times instead of 5. The increased number of repeats was chosen based on preliminary testing, which showed greater variability in performance estimates for TPP models compared to TPC. For parameter optimization, model performance was evaluated using root mean squared error (RMSE). Finally, after the internal CV procedure had determined the optimal parameter, a final model was trained on the full predictor data set.

Separate models were trained to predict each of the 13 harvest traits using data from the control treatment (n_plants_ = 133), drought treatment (n_plants_ = 134), or a pooled dataset containing both treatments (n_plants_ = 267). Each temporal trait at each time point was treated as an independent predictor.

Harvest trait prediction models were trained using all traits measured at all time points (n_predictors_ = 850) and using only measurements from the first half of the experiment (n_predictors_ = 368). Model performance was compared using R² values originating from the repeated internal CV using the final optimal parameter, with statistical significance assessed using either ANOVA followed by multiple pairwise t-tests or a Kruskal-Wallis test followed by multiple pairwise Wilcoxon rank-sum tests, depending on the normality of R² distributions. To correct for multiple testing, Holm’s method was applied to adjust p-values.

Final models were tested on their training data set as well as the other data groups after readjusting them to the fitting z-transformation.

### Variance decomposition of traits

Mixed effects linear models were used to model the temporal and harvest traits. Temporal and harvest traits were modelled similarly, except that the model used for temporal traits, (Eq. 1) included a temporal term, which was not the case for the harvest trait model, (Eq. 2). In addition, for harvest traits, the genetic repeatability (GR) was estimated using the genetic and residual variance components (Eq. 3).

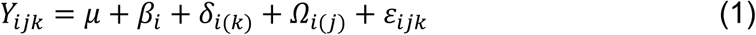

where

|  |  |  |  |
| --- | --- | --- | --- |
| 301 | Y | temporal trait value |  |
| 302 | $\mu$ | overall mean trait value for all plants | |
| 303 | $\beta_i$ | random effect of the ith measuring time point | $\sim N(0, \sigma^2_{\beta_i})$ |
| 304 | $\delta_{i(k)}$ | random effect of the treatment within ith time point | $\sim N(0, \sigma^2_{\delta_{i(k)}})$ |
| 305 | $\Omega_{i(j)}$ | the random effect of jth genetic line within ith time point | $\sim N(0, \sigma^2_{\Omega_{i(j)}})$ |
| 306 | $\varepsilon_{ijk}$ | pooled error of the individual at ith time point | $\sim N(0, \sigma^2_{\varepsilon_{ijk}})$ |
| 307 |  |  |  |

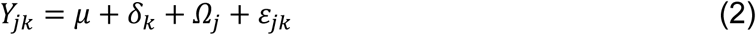

where

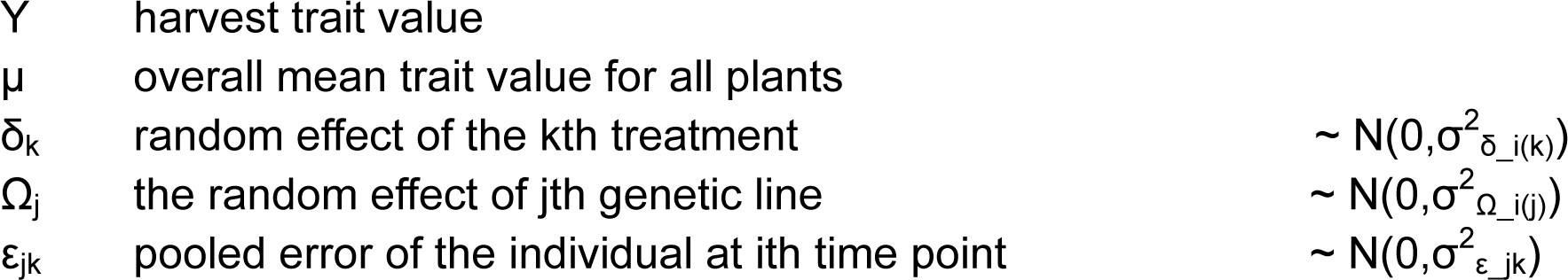

|  |  |  |  |
| --- | --- | --- | --- |
| 310 | Y | harvest trait value |  |
| 311 | $\mu$ | overall mean trait value for all plants | |
| 312 | $\delta_k$ | random effect of the kth treatment | $\sim N(0, \sigma^2_{\delta_{i(k)}})$ |
| 313 | $\Omega_j$ | the random effect of jth genetic line | $\sim N(0, \sigma^2_{\Omega_{i(j)}})$ |
| 314 | $\varepsilon_{jk}$ | pooled error of the individual at ith time point | $\sim N(0, \sigma^2_{\varepsilon_{jk}})$ |
| 315 |  |  |  |

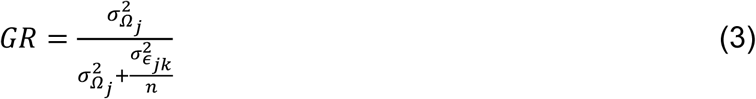

where

|  |  |  |
| --- | --- | --- |
| 318 | GR | The genetic repeatability of a harvest trait |
| 319 | $\sigma^2_{\Omega_j}$ | The genotypic variance |
| 320 | $\sigma^2_{\varepsilon_{jk}}$ | The residual variance |
| 321 | n | The number of replications |
| 322 |  |  |
| 323 |  |  |
| 324 |  |  |

## Results

### Genotypic clustering and drought tolerance ranking of barley lines

To visualize the genotypic and treatment-specific grouping patterns, k-mean clustering (with k = 2) was applied on PCA projected data. The results showed that separation of clusters can be observed based on PC1 (36.6%) and PC2 (31.3%) (Fig. 2A). Notably, among the genetic lines, L6 (from HEB population) clustered separately from the other lines, highlighting potential differences in response to treatment conditions. Moreover, plants grown under control conditions were separated from drought-stressed plants, highlighting that treatment effects contribute to variance in the data.

**Figure 2.**
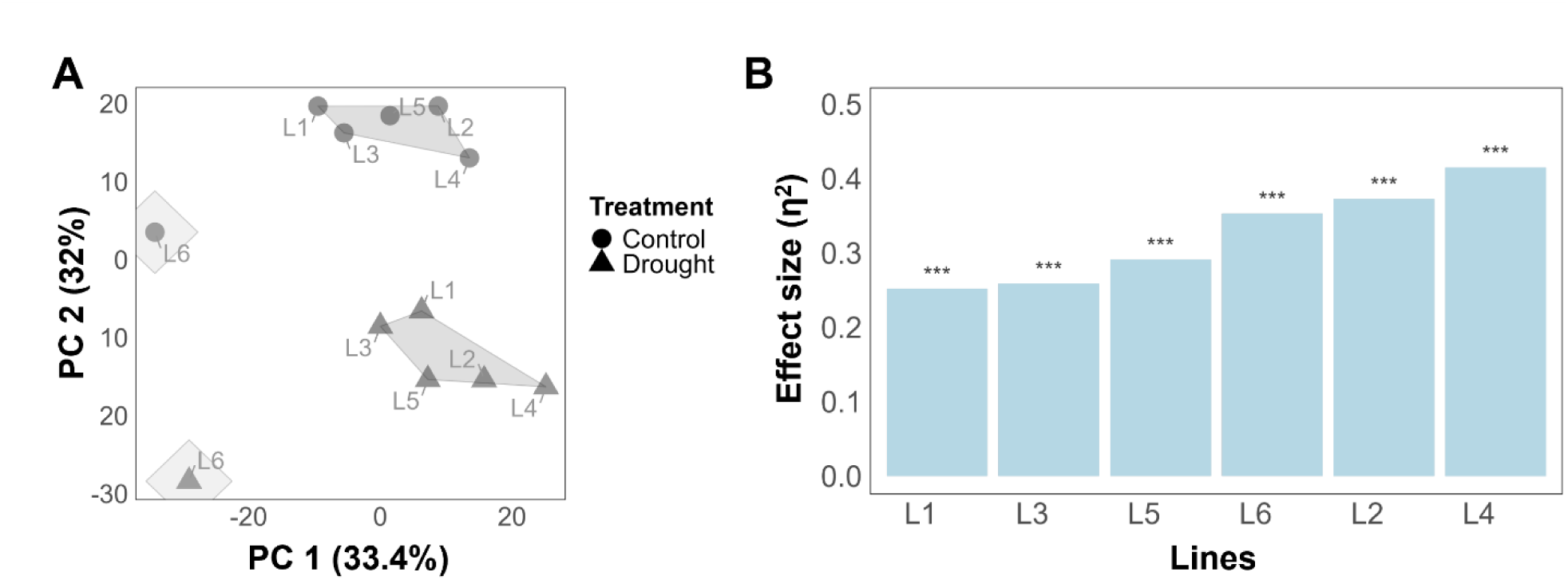
Treatment separation and ranking of the barley lines according to their susceptibility to drought stress. A. k-means clustering using a two-dimensional PCA based on all traits across all time points for the six barley lines. Treatments are represented as control (circle shape) and drought (triangle shape). B. Tolerance ranking of genotype to drought stress using PERMANOVA quantifying significance and effect size of treatment on temporal traits. The asterisks represent a significance level P-value < 0.001.

Tolerance ranking, i.e. quantifying the effect of drought on temporal or harvest traits, found that drought has a highly significant effect (p < 0.001) on temporal traits in all lines (Fig. 2B). By effect size, L1 appeared most tolerant in temporal traits (η^2^ = 0.228), closely followed by L3 (η^2^ = 0.248) and L5 (η^2^ = 0.274). The same three lines appeared most tolerant when harvest traits were analyzed instead of temporal traits (Supp. Fig. S1), with lines L2, L4 and L6 exhibiting the least amount of tolerance with respect to both harvest and temporal traits.

### Temporal phenomic classification of treatment

Temporal phenomic classification (TPC) of plant treatment (drought vs. control) was performed using all temporal predictors from daily time points across the full duration of the experiment (non-aggregated predictors) where model mean accuracy was 0.99 (Fig. 3A). In another approach, predictors were aggregated by week using summary statistics (mean, minimum, and maximum), as described in the methods section, resulting in a slightly lower but maintained high mean accuracy of 0.98 compared to daily time points (Fig. 3B). Variable importance analysis revealed that canopy temperature depression (ΔT) at early-stage (3 weeks of drought stress), along with, as expected, RGB-based plant size estimates (area from RGB side view and plant volume) at late-stage (longer duration of drought stress) were the most influential predictors of treatment classification, regardless of daily or weekly aggregated data (Fig. 3 and Supp. Table S6).

**Figure 3.**
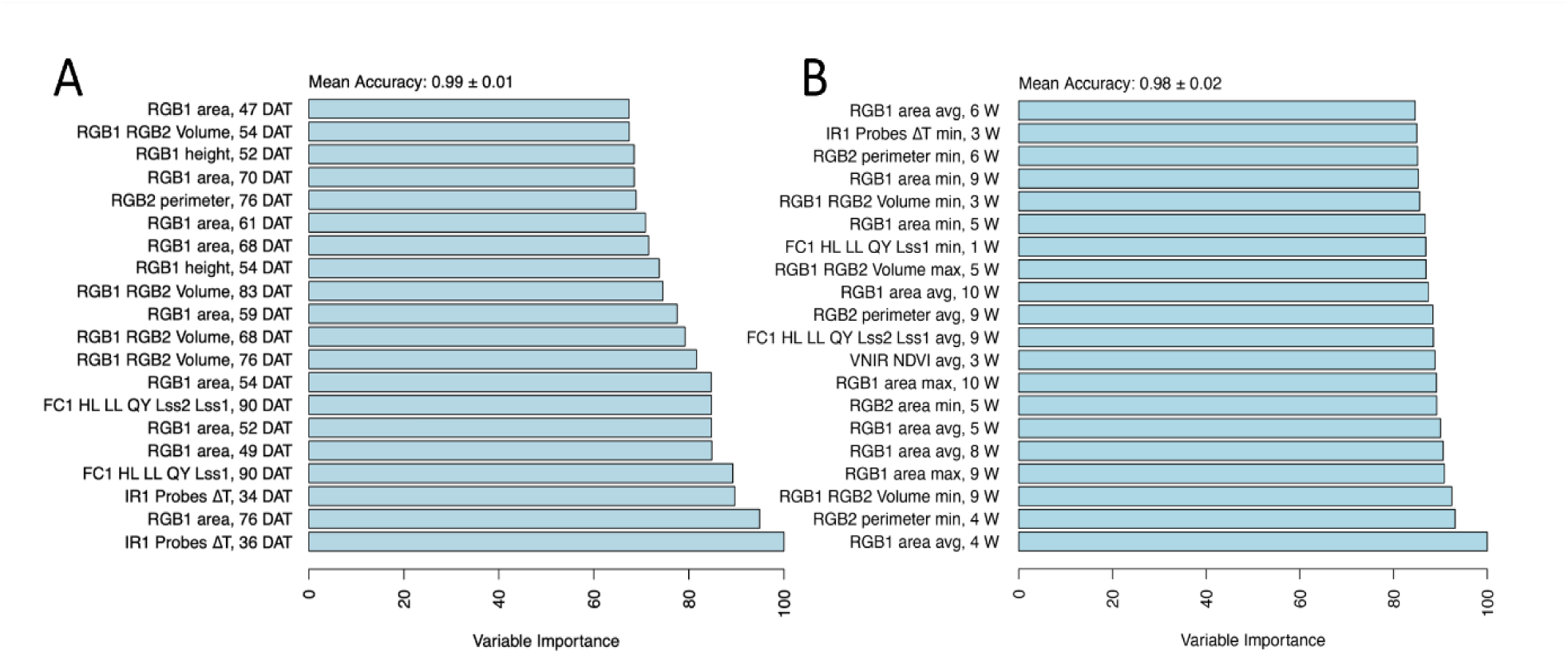
Variable importance for classification of treatment. The importance of the temporal traits was assessed using a random forest model with a mean accuracy metric for A. non-aggregated data set and B. aggregated data set composed of minimum, maximum, and average per week, and top 20 traits differentiating between treatments. Variable importance was determined using a permutation-based method, where the model’s prediction accuracy was compared before and after shuffling each variable. Higher importance values indicate a greater decrease in accuracy when a variable is permuted, signifying its stronger contribution to classification performance.

In addition, TPC using daily (non-aggregated) predictors was also performed by training separate models using only the data from individual weeks. All week-specific models, except the one based solely on week 0 (during which the drought treatment was initiated), achieved high accuracy in distinguishing treatments (0.973 ≤ mean accuracy ≤ 0.99; Supp. Fig. S2). The model trained exclusively on week 0 data had lower performance (mean accuracy = 0.695), likely due to the limited physiological response at this early stage. Notably, classification models using only the second and third weeks of measurements (corresponding to week 1 and 2 after inducing drought stress) relied almost entirely on ΔT estimates, along with traits from chlorophyll fluorescence and visible-near-infrared (VNIR) imaging (Supp. Fig. S3). This finding underscores the importance of these traits in early drought stress detection.

By comparing predictors derived from the daily and weekly-aggregated data, we found that both contributed to identifying critical time points in the stress response, where common traits at specific time points were identified. Notably, from the chlorophyll fluorescence (morning protocol), the ratio of quantum yield under low to high light (FC1 HL LL QY Lss2/Lss1) was selected at day 90 and week 9, reflecting the severe impact of stress responses at the late stage (Fig. 3).

### Temporal phenomic prediction of harvest traits

Temporal phenomic prediction (TPP) was also performed for all harvest traits using RF regression and LASSO models, with training data from the control treatment, drought treatment, or the pooled data set containing both treatments. Among the predicted traits, total biomass dry weight, spike number, total spike weight, and five spikes weight were generally the most predictable based on R² values (Fig. 4A). Therefore, our subsequent comparisons of prediction accuracy focus on these four traits. Among these traits, total biomass exhibited substantially higher predictability compared to the other traits.

**Figure 4.**
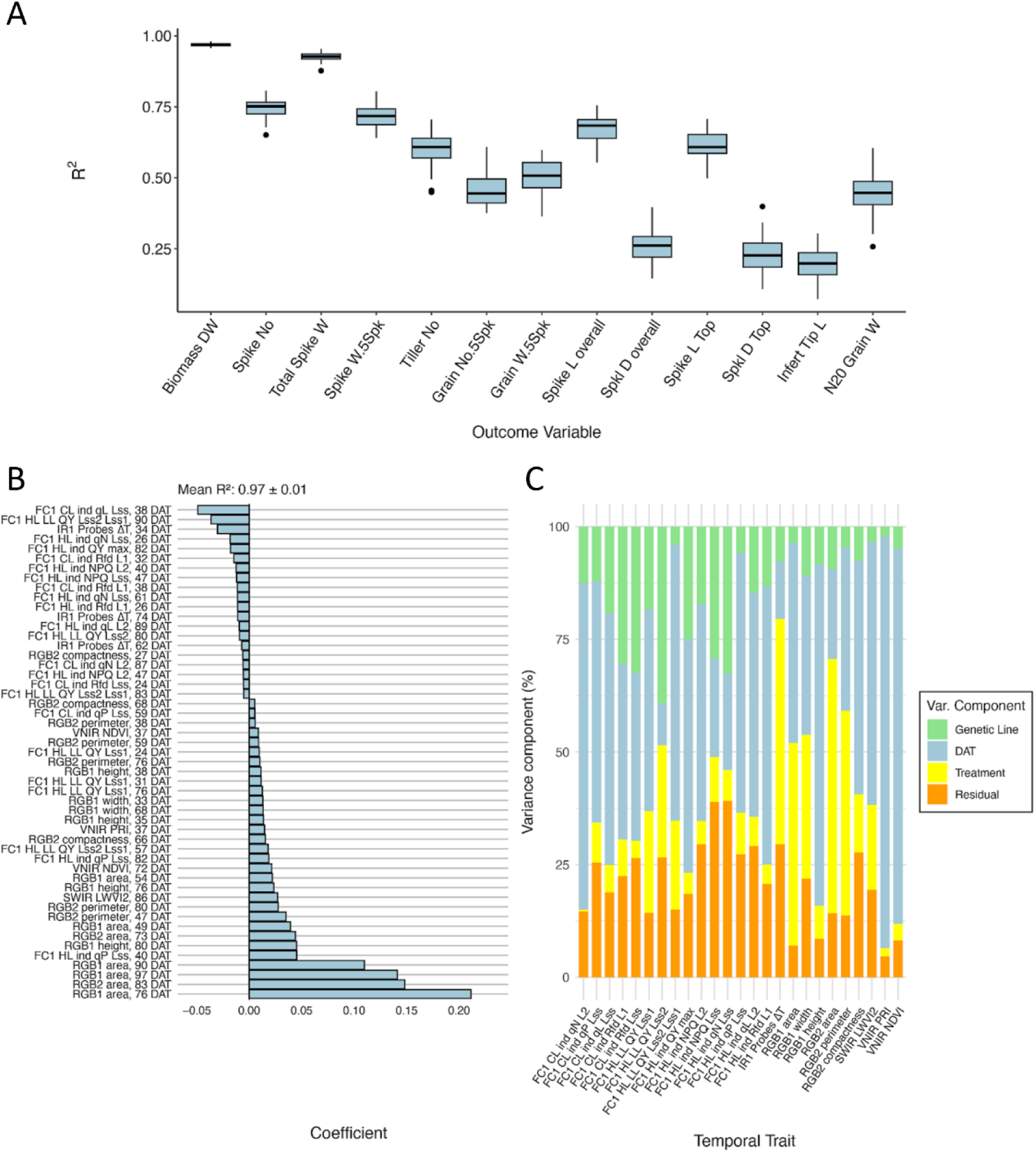
Performance of the prediction models for harvest traits. A. Boxplots showing the accuracy (determined by R^2^) among all the harvested traits using a LASSO model trained on pooled non-aggregated dataset. B. LASSO coefficient on the total biomass dry weight. Coefficients with absolute values below 0.005 are not plotted C. Partitioning of variances of the temporal traits selected from LASSO model.

The relative performance of RF compared to LASSO depended on the training data set (i.e., control, drought, or the pooled treatment data) and the response trait, with only minor differences in overall accuracy between the two models (Supp. Fig. S4). Due to the small and unsystematic difference in performance between these model types, the rest of the findings reported will focus on the LASSO models, as they are facile to interpret without additional feature importance scoring. The choice of training data set (i.e., drought, control, or pooled) had a more pronounced effect on model performance than the choice of modelling approach. Models trained on the pooled dataset generally performed best, followed by those trained on drought data (Supp. Fig. S5). There is a slight overlap between useful predictors in pooled data TPP and TPC of treatment, suggesting that these models, by implicitly capturing treatment-related variance, leveraged the drought-induced variability in both predictors and response traits to improve robustness (e.g., TPC, Fig. 3A and biomass prediction, Fig. 4B). This finding is further supported by the fact that many harvest traits, especially total biomass, spike weight and spike number, showed a large variance contribution from treatment (Supp. Fig. S6). However, when using the pooled model to predict harvest traits using only drought or control data, the performance was marginally affected (Supp. Table S7), indicating a reasonable degree of transferability of the model.

LASSO coefficients for pooled and drought treatment biomass prediction were generally associated with plant size estimates (e.g., plant area) derived from RGB imaging at late time points (Fig. 4B). These predictors not only had the largest absolute coefficient values but also showed high variance contributions linked to the treatment (Fig. 4B-C and Supp. Fig. S7). Aggregating predictors by week caused a slight drop in model performance during TPP, but accuracies were still similar to those found in the original daily data set (Supp. Fig. S8).

To assess the feasibility of early prediction of harvest traits and identify early predictors of importance, models were retrained using only measurements from the first half of the experiment (Supp. Fig. S9). Compared to models trained on all time points, R² prediction accuracy was decreased and had higher variance in all models. However, despite this reduction in accuracy, predictions remained reasonably effective, particularly for biomass dry weight (mean R^2^ 0.924 for pooled-treatment LASSO model) and total spike weight (mean R^2^ 0.837 for pooled-treatment LASSO model) (Supp. Fig. S10). In these early-timepoint models, pooled-treatment models still resulted in the highest overall accuracy. RGB plant size estimates were again found to be important predictors, but they were supplemented with predictors from other sensors. Biomass prediction specifically included hyperspectral indices (MCARI1, LWVI2, NDVI) while total spike weight more strongly relied on photosynthetic efficiency from chlorophyll fluorescence and also included plant senescence estimation by RGB color indices at 49 DAT and ΔT from thermal IR imaging (Supp. Fig. S10).

## Discussion

Our study investigated and demonstrated the potential to leverage temporal high-throughput phenotyping in dissecting plant responses to drought stress and in predicting agronomically relevant outcomes in different barley lines. By gathering and analyzing a comprehensive set of temporal phenotypic traits measured across the growth cycle, we developed models capable of accurately classifying drought treatment and predicting harvest-related traits such as biomass dry weight, spike weight, and spike number. Notably, treatment classification achieved high accuracy (mean accuracy ≥ 0.97) when using temporal predictors from any week except the first, during which drought treatment had only recently commenced.

### Model accuracy and cross validation

An important outcome of the harvest trait prediction analysis was the clear superiority of models trained on the pooled dataset. This increase in accuracy was likely due to structured variability in temporal and harvest traits introduced by treatment, as can be seen in the substantial treatment-induced variance in predictors and response traits, as well as in the overlap in important predictors from TPC and pooled treatment TPP. In addition to achieving the highest accuracy across internal cross-validation folds, pooled treatment models also performed well when tested separately on drought and control treatment data (Supp. Table S7). While this test does not constitute a true external cross-validation, it provides evidence that the improved performance of pooled models is not merely driven by treatment-level separation. The high accuracy observed when predicting within each treatment group suggests that the models capture meaningful, continuous variation among individual plants, rather than just categorical differences between drought and control conditions.

External validation, in which a model is tested on data entirely excluded from training, remains the most rigorous method for identifying overfitting and assessing model generalization. While external validation was not feasible in this study, model performance was evaluated using repeated internal cross-validation, where R² values were computed on data withheld from training. This approach reduces the risk of overfitting. Apart from this, other observations are indicative that overfitting was not the cause for the high observed accuracy.

Most notably, prediction accuracy varied substantially across response traits despite the use of identical predictor sets. Traits with limited or indirect association to vegetative growth dynamics, such as infertile tip length and spikelet density, consistently yielded low predictive performance (R² < 0.3). If the models were overfitted, one would expect uniformly high accuracy across traits, regardless of biological relevance. Instead, this pattern suggests that the high accuracy observed for traits, such as biomass and spike weight, likely reflects a reliable predictive signal rather than a model artefact. A similar argument holds for the treatment classification task, while most weekly models achieved near-perfect accuracy, classification using only the first week of measurements performed poorly. This was biologically expected, as the drought treatment had only just started during this week (WP 0) and had not yet induced measurable phenotypic changes. The presence of both high- and low-performing models, in line with biological expectations, provides indirect evidence that overfitting is not the primary driver of the observed model performance.

Moreover, both regression methods used LASSO and Random Forest have inherent mechanisms for mitigating overfitting. LASSO applies L₁ regularization, which penalizes model complexity by shrinking coefficients and effectively selecting a subset of informative features (Ying, 2019). Random Forests, through ensemble learning and the use of out-of-bag error estimation, reduce the risk of overfitting by averaging across multiple decorrelated decision trees (Ghojogh & Crowley, 2023). Nonetheless, while these approaches help control overfitting, a more reliable way to assess generalizability is through external cross-validation using independent data not seen during model training or optimization (Ghojogh & Crowley, 2023). Due to the limited sample size and experimental design, such validation was not feasible in this study, but should be prioritized in future work.

### Integrative insights into model interpretability, classification, and prediction

Variable importance was assessed using model coefficients in LASSO and a permutation-based approach in Random Forests. These measures provide insight into predictive relevance, but reflect conditional importance, may fail to identify all relevant predictors when multicollinearity is present, as only one among a set of correlated variables may be selected or appear important (Grömping, 2009). This is likely the reason for the slight inconsistencies observed in variable importance rankings between models using daily (non-aggregated) and weekly (aggregated) predictors.

#### Temporal phenomic classification of treatment

For treatment classification, canopy temperature depression (ΔT) at early time points and RGB plant size estimates at late time points emerged as important predictors (Fig. 3). Notably, treatment could be classified very accurately early in the experiment mainly by relying on ΔT measurements, highlighting the importance of this trait for early drought detection. Plant canopy temperature has an impact on plant growth by non-linearly regulating photosynthesis, respiration, and transpiration rates. By increasing water deficit, the efficiency of cooling the leaf surface through transpiration diminishes, leading to an increase in leaf temperature (Biju et al., 2018; Way & Yamori, 2014). Therefore, changes in canopy temperature provide a valuable proxy of stomatal regulation and an indicator of different stress responses (Wen et al., 2023). Stomatal closure serves as an initial response to drought stress to prevent excessive water loss, leading to alterations in physiological response and metabolic pathways (Farooq et al., 2024). Thus, a rise in canopy temperature was observed in higher ΔT in drought-stressed plants. However, as drought duration increases over time, ΔT became a less significant classifier of treatment, likely due to physiological adaptations such as stomatal acclimation, osmotic regulation, and changes in transpiration dynamics, which can moderate canopy temperature despite continued water limitation (Lawson & Blatt, 2014). In addition, plants progressively reduce their biomass due to water limitations (Pham et al., 2019). This structural change, including leaf area reduction, alters canopy-atmosphere interactions and subsequently affects canopy temperature regulation (Vico et al., 2023).

#### Temporal phenomic prediction of harvest traits

For the prediction of harvest traits, RGB-based plant size estimates from late time points emerged as dominant predictors, particularly in models for biomass (Fig. 4). This is likely due to the strong biological alignment between total biomass and plant size, as both reflect cumulative vegetative growth. This aligns with previous studies showing the reduction of biomass accumulation under drought stress (Cai et al., 2020; Neumann et al., 2015). Moreover, the high correlation between the biomass and yield, where biomass reduction ultimately affects yield-related traits, as a result of low assimilates for grain production, reflecting a source-to-sink limitation (Findurová et al., 2023; Rosati & Benincasa, 2023). When prediction models for biomass and total spike weight were trained using only pooled-treatment predictors from the first half of the experiment, RGB plant size traits remained influential, but they were complemented by features derived from all of the other sensors. Despite a significant reduction in predictive accuracy compared to models trained on the full dataset, these early-timepoint models still achieved strong performance, with mean R² values of 0.92 for biomass and 0.84 for spike weight, respectively (Supp. Fig. S9 and S10). The relatively high accuracy of these early TPP models is particularly promising in a breeding context, as it demonstrates that complex cumulative traits can be reliably predicted well before they are phenotypically expressed. Harnessing this predictive capacity would allow breeders to select plants early in the breeding pipeline, significantly reducing cost.

Notably, the fraction of open Photosystem II (PSII) centers at light steady state (qL_Lss) showed importance in LASSO coefficient when prediction models were trained on pooled and drought data during the early drought stage (stem elongation stage) (Fig. 4 and Supp. Fig. S7 and S8). The variation in the opening of the PSII centers is probably reflecting alterations in photosynthetic efficiency and electron transport pathways, which shows major mitigation mechanisms to alleviate the negative effects of moderate drought stress (Qiao et al., 2024; Shin et al., 2021). By increasing the stress intensity and duration, the quantum yield of PSII (QY Lss2/Lss1) was observed at the late stage as important predictor, indicating impairment of PSII function, where plants were unable to efficiently transfer energy from high to low light (Zhou et al., 2019).

#### Implications for breeding and timely selection

One of the primary applications of TPP is enabling the early selection of plants in breeding. In this study, it was demonstrated that this task is feasible with high accuracy for a variety of traits. Even predictors from the first half of the phenotyping period in the models achieved high accuracy for traits such as total biomass and total spike weight, potentially allowing selection after only a few weeks of vegetative growth. However, it should also be noted that models trained on predictors from the full phenotyping period remain valuable to breeders, as phenotyping concluded 31 days before harvest. Nevertheless, daily phenotyping and harvesting are both costly and time-consuming processes, which could be partially avoided by implementing early phenotyping to identify tolerant plants prior to the later stages of evaluation (Adak et al., 2023).

An increase in prediction accuracy was observed when data from both drought and control treatments were combined in the training set. This illustrates a key advantage of phenomic prediction over genomic prediction. While genomic prediction captures only the static genetic contribution to trait variation, phenomic prediction leverages genetic, environmental, and genotype-by-environment interaction effects, as phenotypic traits, both predictors and responses, reflect the integrated output of all these factors (Adak et al., 2023). Variance decomposition of both predictor and harvest traits in this study further illustrates that trait variation arises from a combination of genetic background, environmental conditions, and temporal dynamics. In contrast, genomic prediction models are inherently limited to the genetic component of variance and cannot account for time-dependent or environmentally induced effects. The inclusion of a temporal dimension in phenomic prediction further enhanced model performance by capturing dynamic shifts in environmental conditions and their interactions with genotype over time. This ability to model temporal trajectories of plant development and stress response adds substantial predictive power and highlights the unique potential of time-resolved phenomic data in breeding applications.

### Limitations and outlook

Viewed in a breeding context, a limitation of this study is the absence of direct grain yield measurements, which are typically the primary selection targets in breeding programs. While yield-related traits such as total spike weight, spike number, and biomass provide meaningful proxies, they do not fully capture grain production in the field. Nevertheless, the study offers valuable insights into the predictive capacity of temporal phenomic data, and demonstrates how yield-associated traits can be modelled early in the growth period. The approaches presented here are readily transferable to datasets including grain yield, and thus remain relevant for informing breeding strategies.

An additional limitation, as previously discussed, is the absence of external cross-validation for performance assessment in temporal phenomic prediction. Instead, models were evaluated using a repeated 3-fold internal cross-validation procedure, which provides a reliable estimate of predictive performance with minimal overfitting risk, given the available sample size. However, external cross-validation would offer a more robust measure of model generalizability and could be used to assess performance across different environmental conditions. Cross-environment validation has previously been applied in TPP studies to demonstrate its advantage over genomic prediction, which often does not transfer to new environments without a significant decrease in accuracy (Adak et al., 2023; Jarquin et al., 2021). Incorporating genomic prediction alongside the phenomic methods used in this study would also enable a direct comparison of their relative predictive power. However, this strategy would require a substantial increase in the phenotyped population and application of recently developed data integrative approaches (Hobby et al., 2025).

## Acknowledgment

This study in the framework of the CAPILTALISE project was funded by the European Union’s Horizon 2020 research and innovation programme (grant no. 862201). We acknowledge David Hobby for his feedback during the development of the analysis pipeline. We thank Pavla Homolová for helping prepare plant material during the experiment at PSI Research Center, Czech Republic.

## Authors contribution

Conceptualization: AKS, EF, ZN, KP, Methodology & Investigation: LA, BP, AKS, KP, Data Curation & Formal Analysis: HT, LA, BP, Funding Acquisition & Supervision: EF, ZN, KP, Writing – Original Draft: HT, LA, Writing – Review & Editing: All authors.

## Data Availability Statement

Data used in this study are available in the supplementary Files S1-S14.

## Supplementary figures

**Supp. Figure S1.**
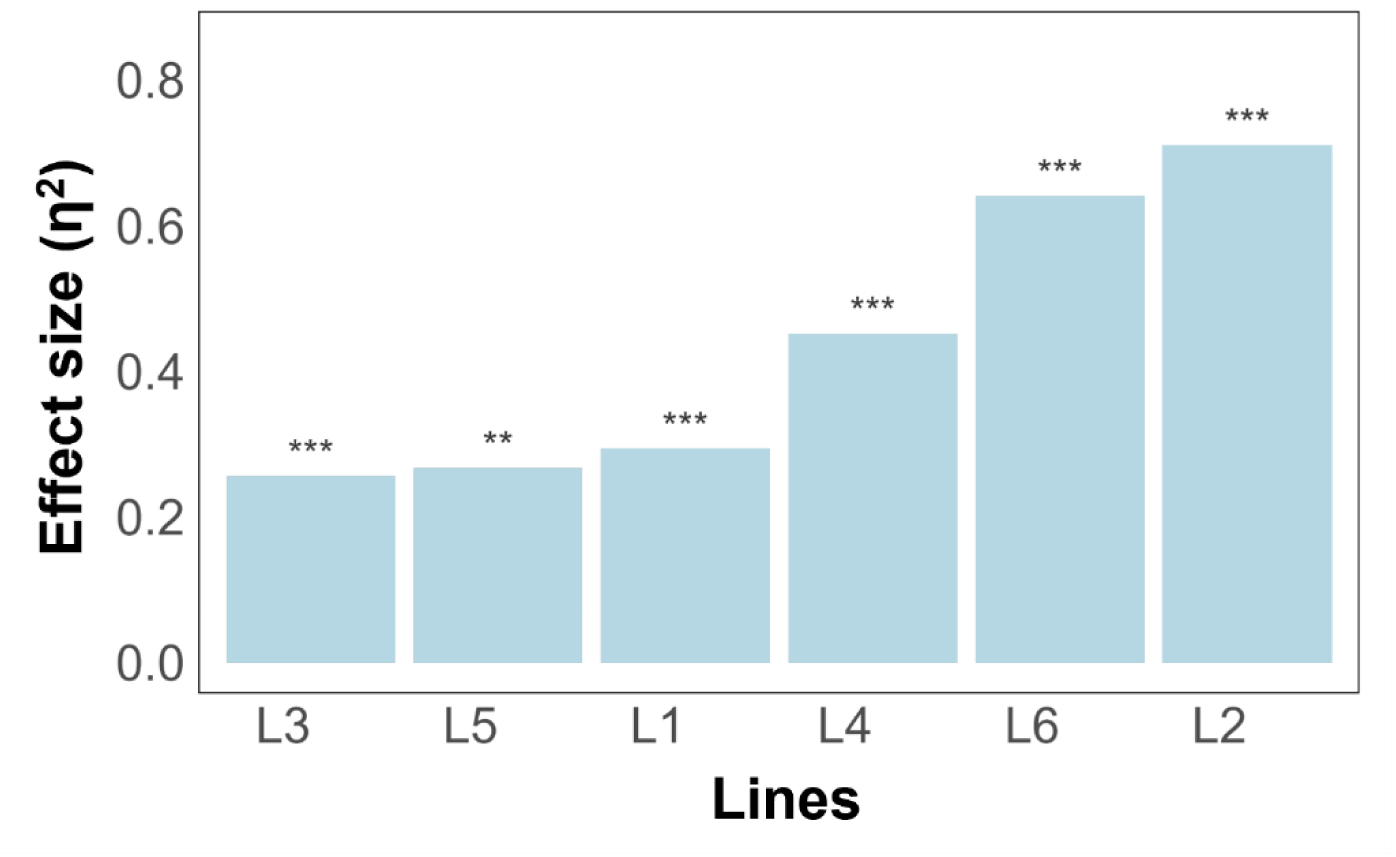
Ranking the lines according to their susceptibility to drought stress on harvest traits. PERMANOVA was used to quantify the significance and effect size of the treatment on harvest traits. The asterisks represent significance level P-value < 0.001 for *** and 0.01 for **.

**Supp. Figure S2.**
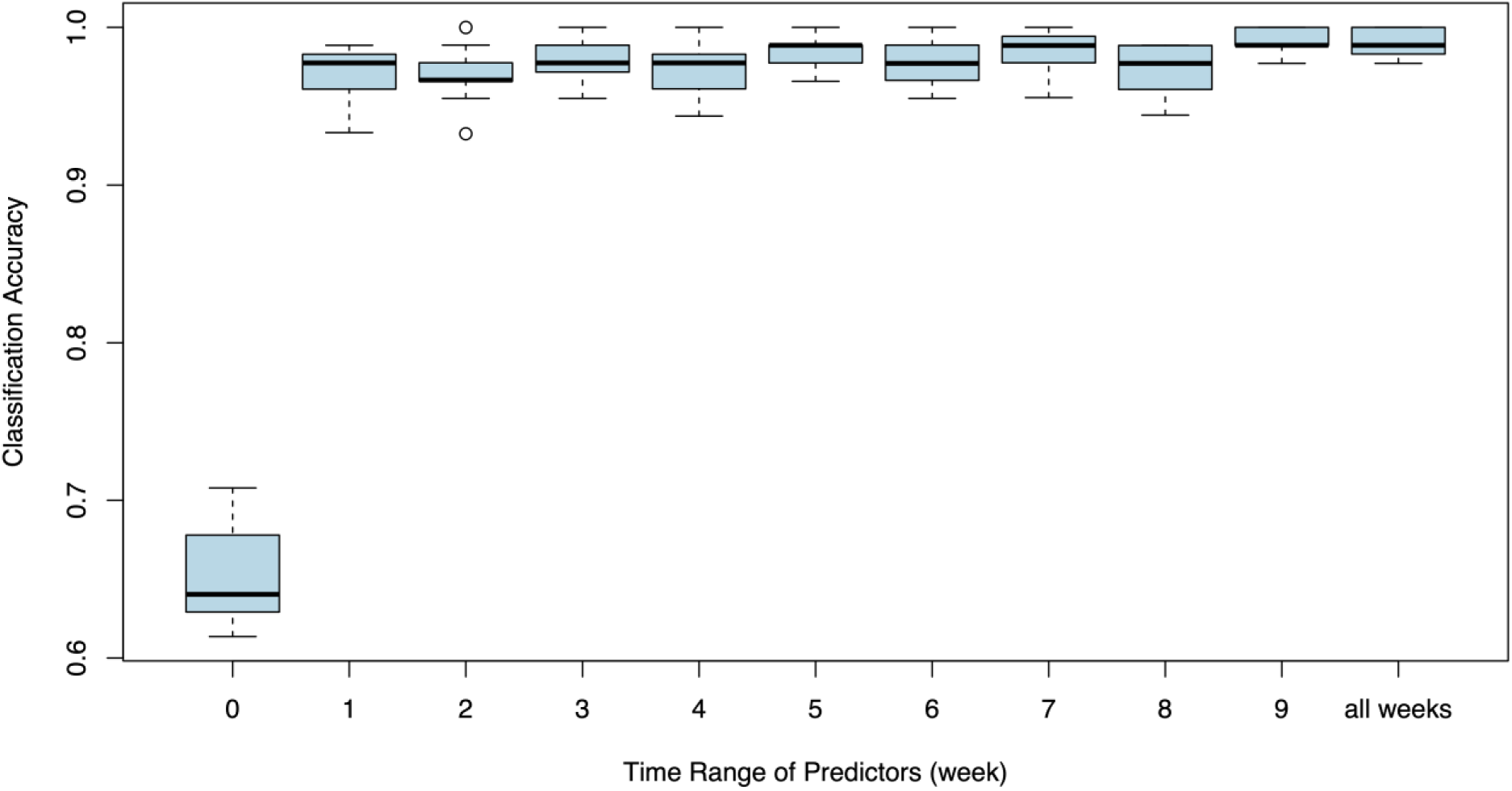
Temporal phenomic classification (TPC) of treatment (drought / control) using predictors subset by week. TPC was performed using random forest models. Predictors from each week were used separately along with a model trained on the full data set (all weeks).

**Supp. Figure S3.**
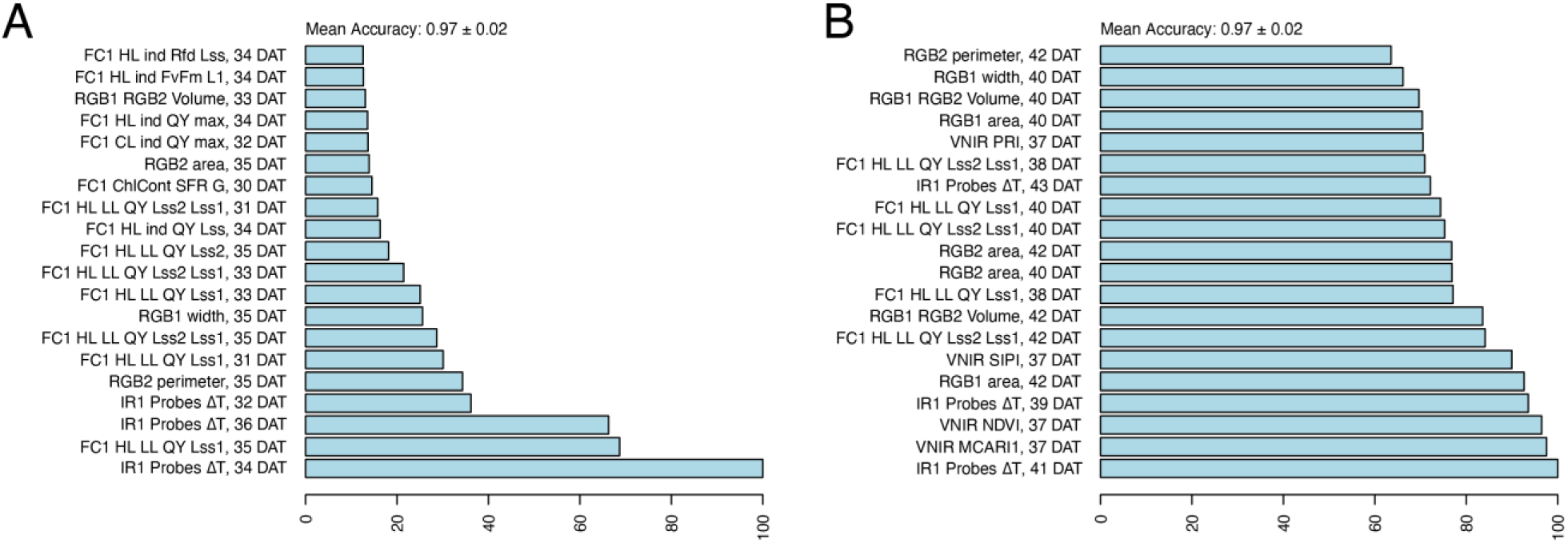
Variable importance of temporal phenomic classification (TPC) of treatment using predictors from two weeks. A. Traits from week 1 and B. week 2 after inducing drought stress. TPC was performed using random forest models.

**Supp. Figure S4.**
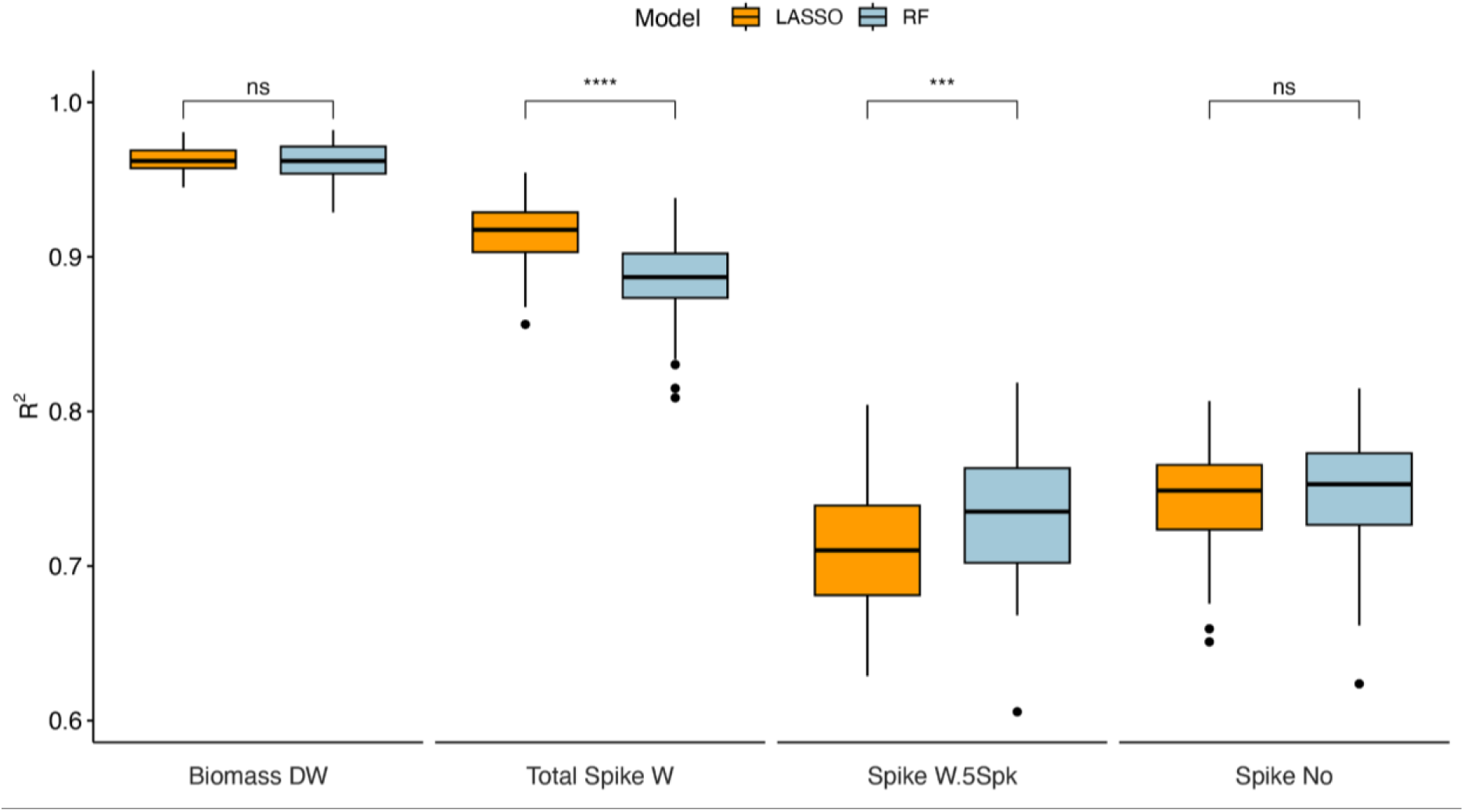
Comparison between the accuracy of models. Boxplots showing the accuracy (determined by R^2^) among selected harvested traits using LASSO and Random Forest models trained on a pooled non-aggregated dataset. The significance level was determined as *** for P < 0.001, **** for P < 0.0001 and ns for non-significant differences between the models.

**Supp. Figure S5.**
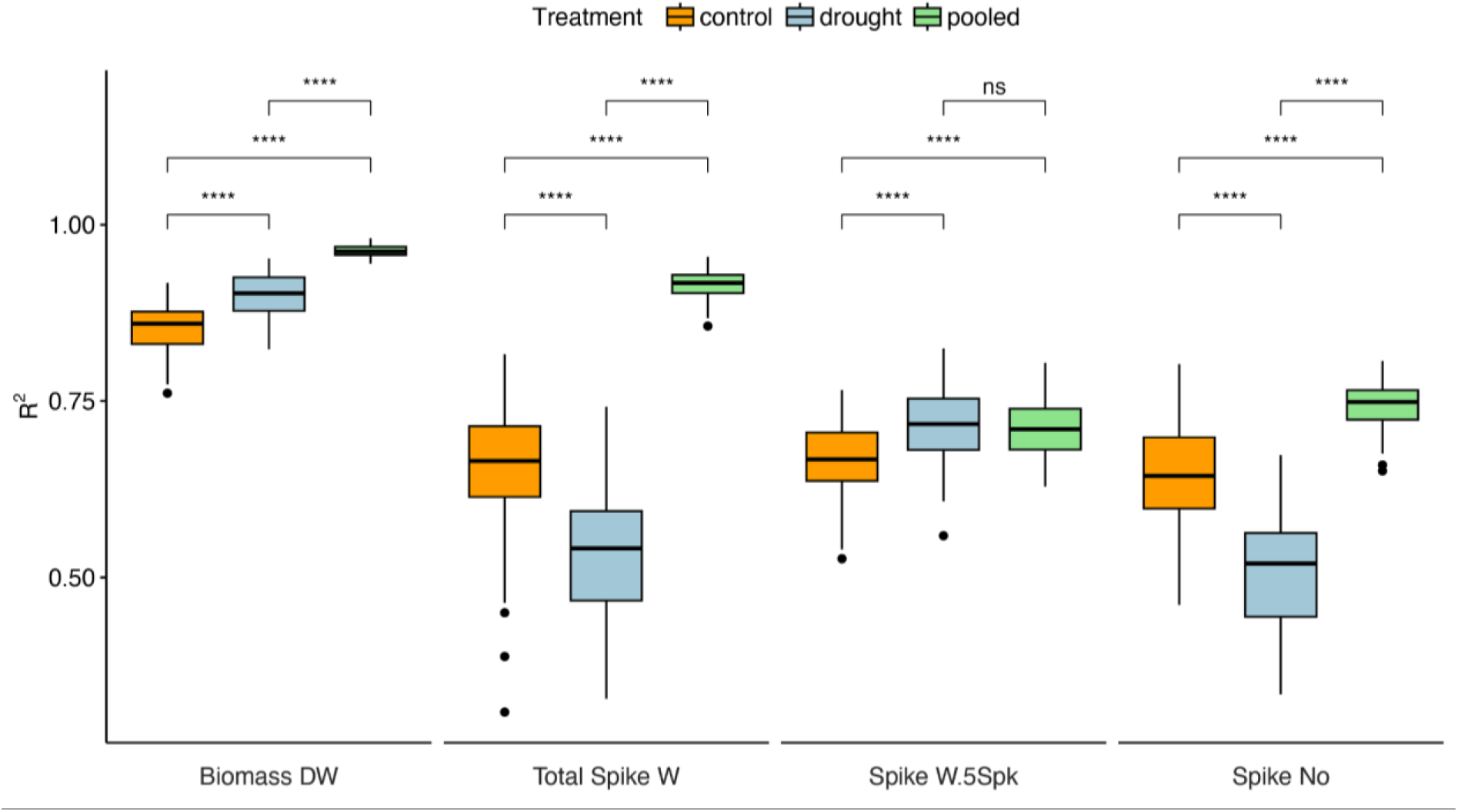
Comparison between the different treatments using the LASSO model. Boxplots showing the accuracy (determined by R^2^) among selected harvested traits using the LASSO model trained on a non-aggregated dataset. The significance level was determined as **** for P < 0.0001 and ns for non-significant differences between the models.

**Supp. Figure S6.**
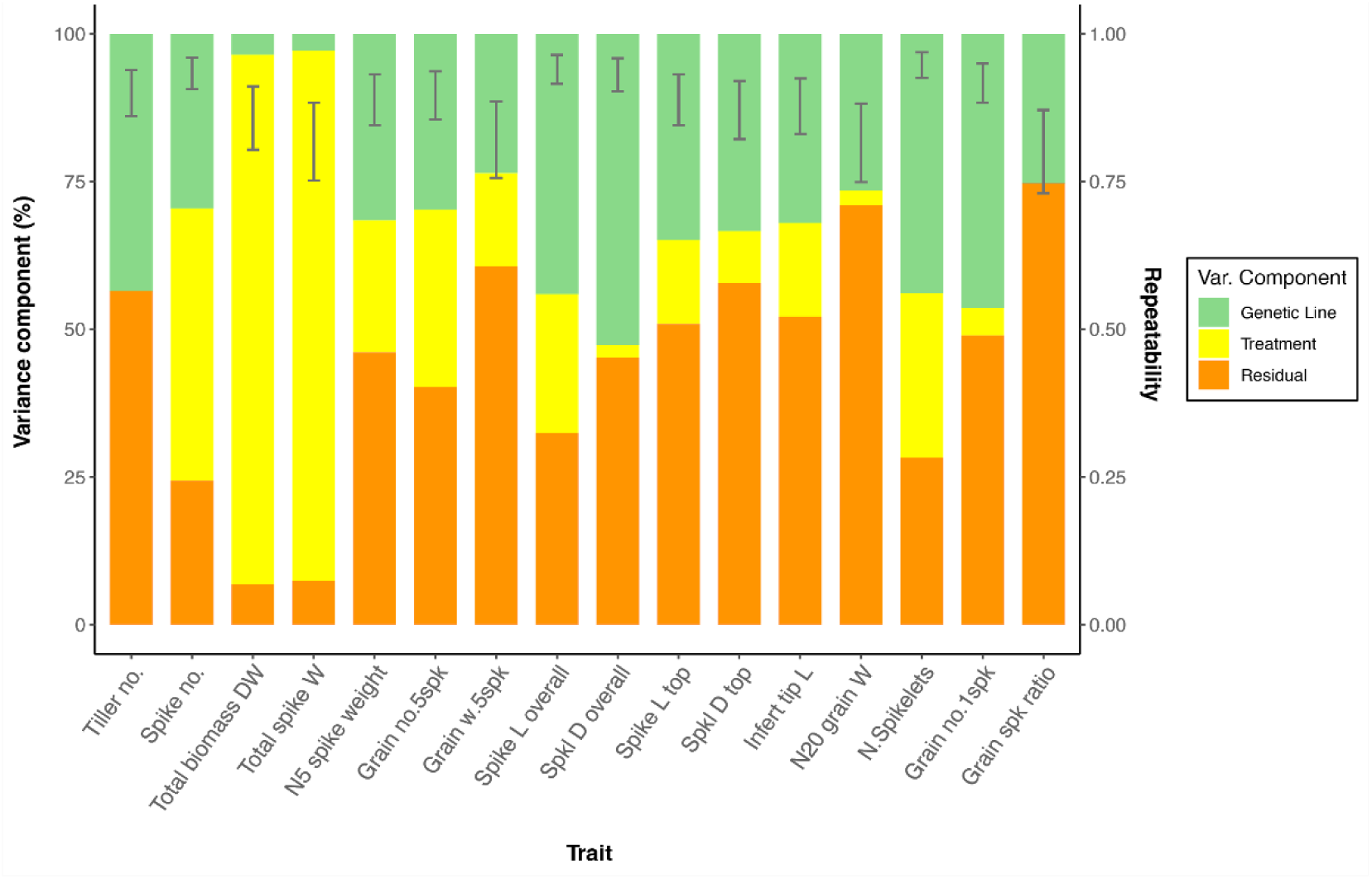
Partitioning of variances and repeatability of the harvest-related traits. Repeatability on the secondary y-axis is represented as error bars.

**Supp. Figure S7.**
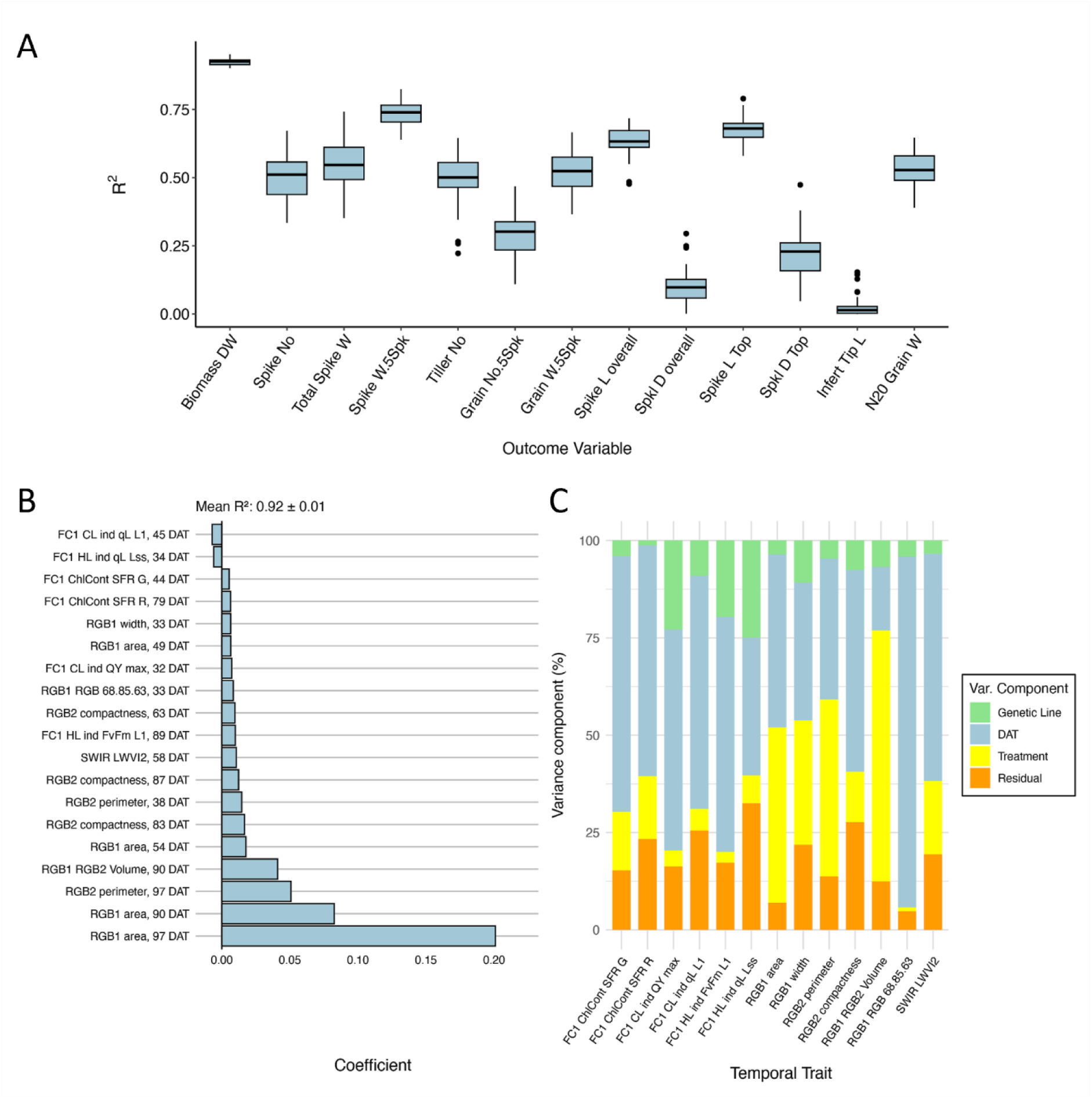
Harvest trait prediction models on drought non-aggregated dataset. A. Boxplots showing the accuracy (determined by R^2^) among all the harvested traits using the LASSO model trained on drought non-aggregated dataset. B. LASSO coefficients on the total biomass dry weight where coefficients with absolute values below 0.005 are not plotted. C. Partitioning of variances of the temporal traits selected from LASSO model.

**Supp. Figure S8.**
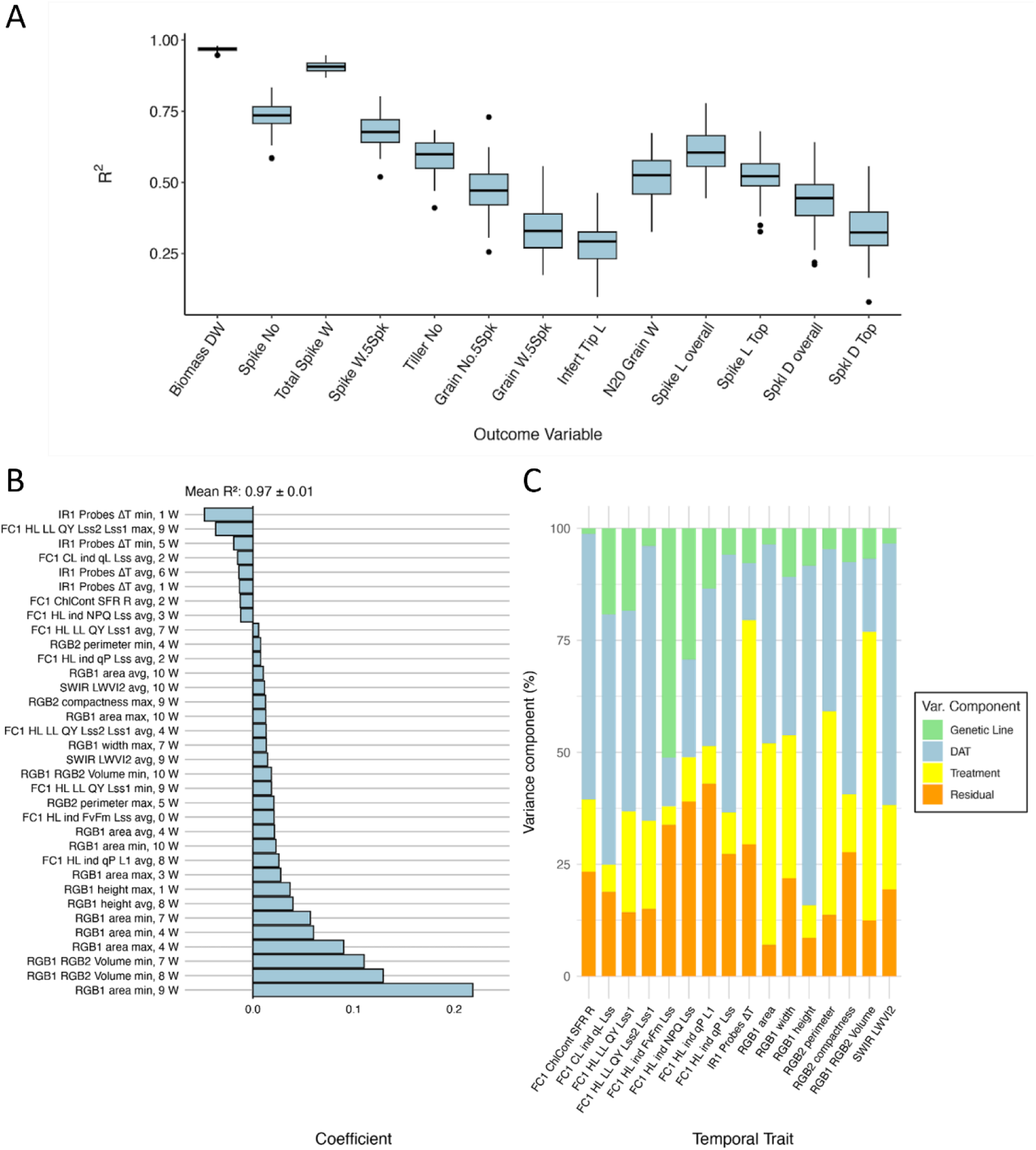
Harvest-traits prediction models on an aggregated dataset. A. Boxplots showing the accuracy (determined by R^2^) among all the harvested traits using the LASSO model trained on pooled aggregated dataset. B. LASSO coefficient on the total biomass dry weight. C. Partitioning of variances of the temporal traits selected from LASSO model.

**Supp. Figure S9.**
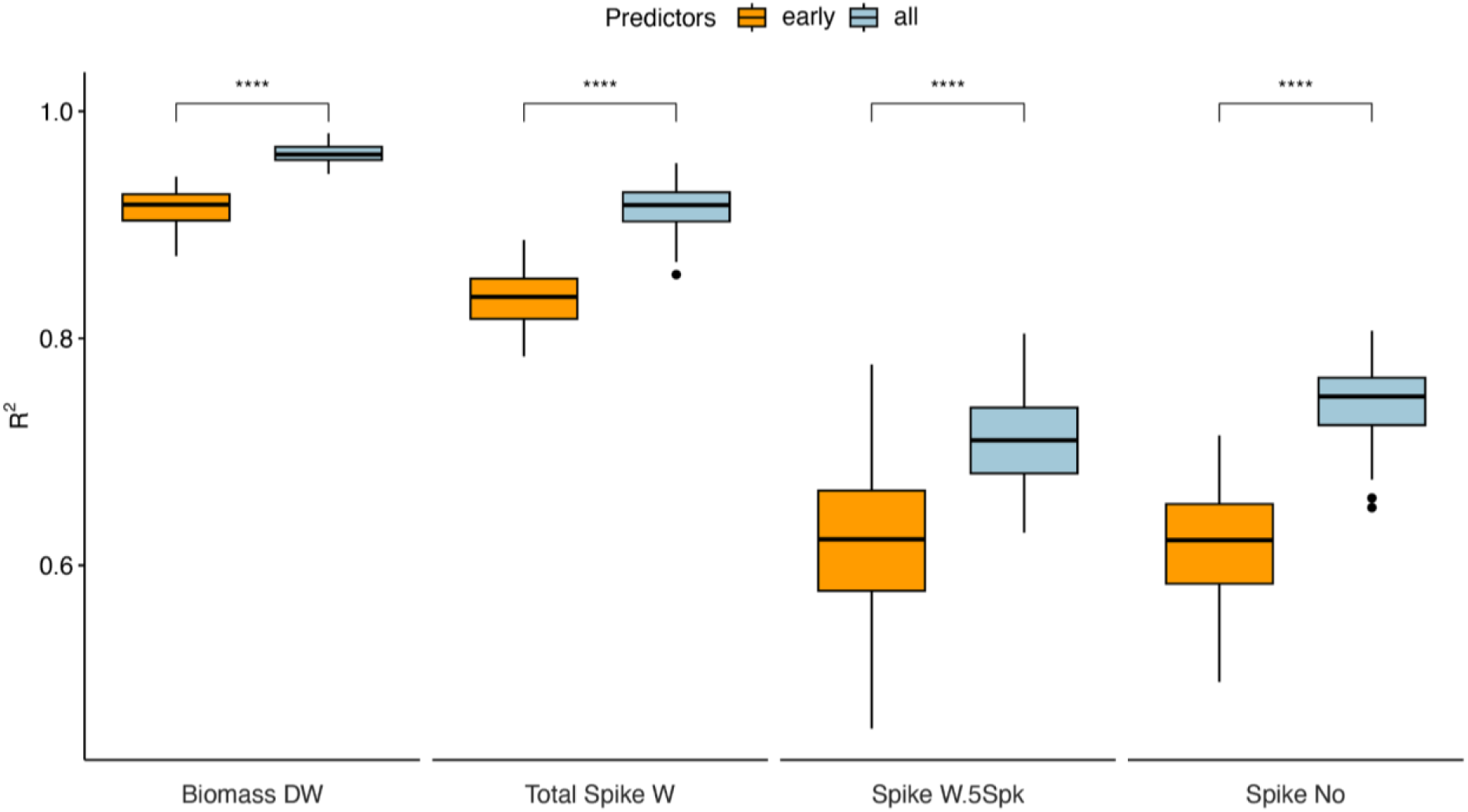
Comparison between the different early and all time points using the LASSO model. Boxplots showing the accuracy (determined by R^2^) among selected harvested traits using the LASSO model trained on pooled non-aggregated dataset. Early time points were selected until 51 days after transplant (DAT). The significance level was determined as **** for P < 0.0001 and ns for non-significant differences between the models.

**Supp. Figure S10.**
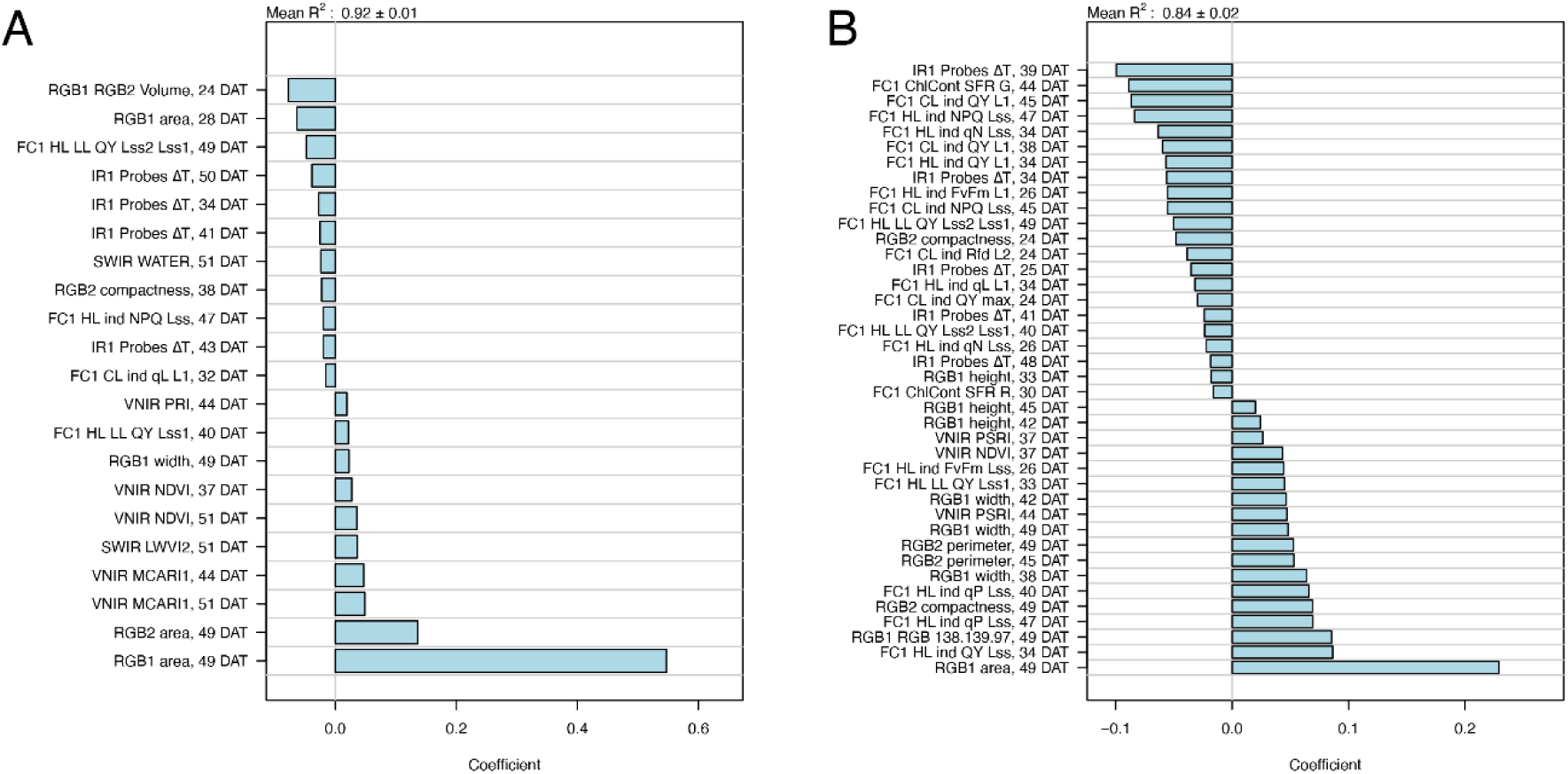
LASSO coefficients of early time point TPP models for harvest traits. A. Biomass dry weight and B. total spike weight were selected. Only coefficients with absolute values above 0.015 are plotted.

## Supplementary tables

**Supp. Table S1.**
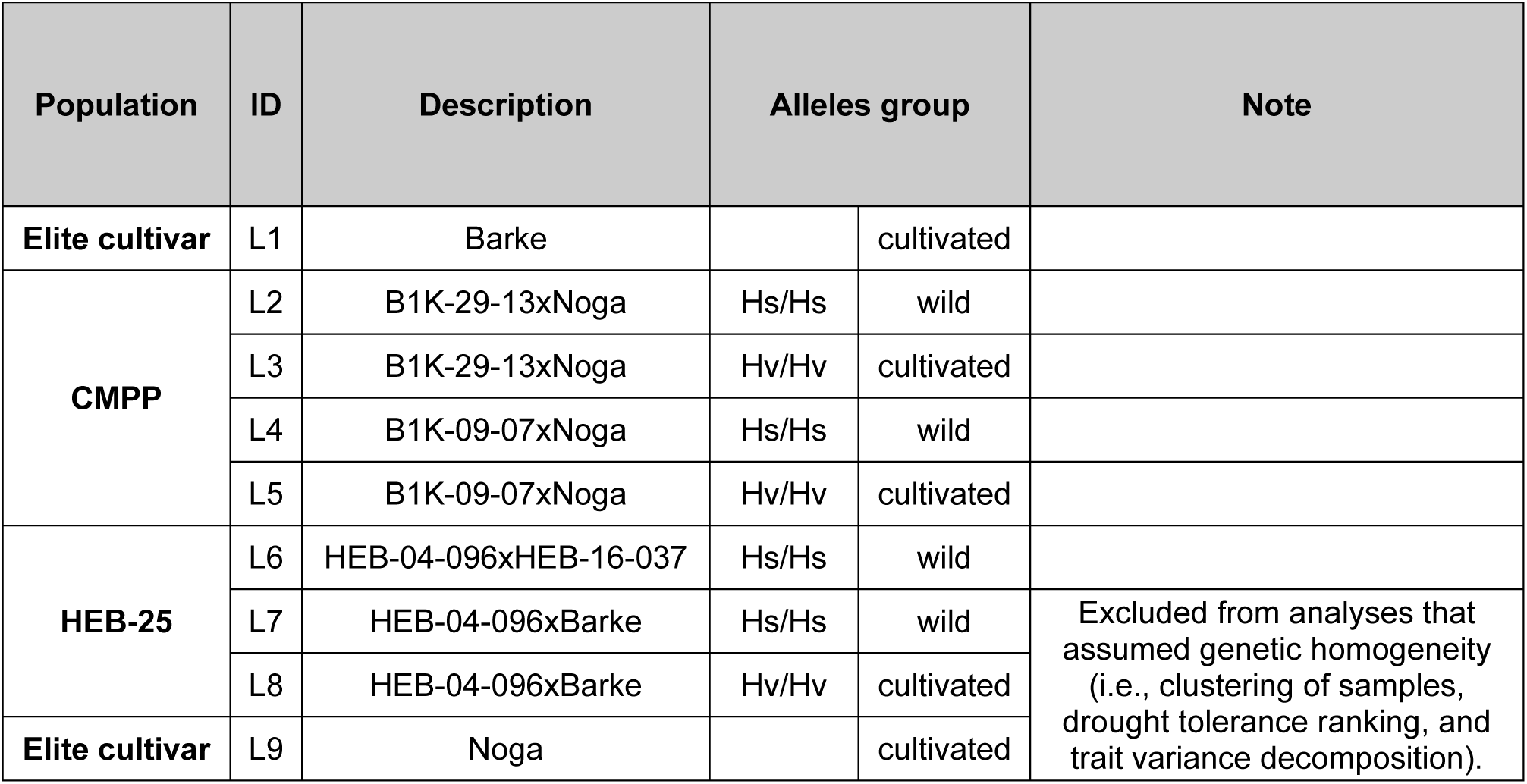
Genetic background of the plant material for the selected lines.

**Supp. Table S2.** List of temporal traits description from multiple imaging sensors and harvest-related traits. (Excel file)

**Supp. Table S3:**
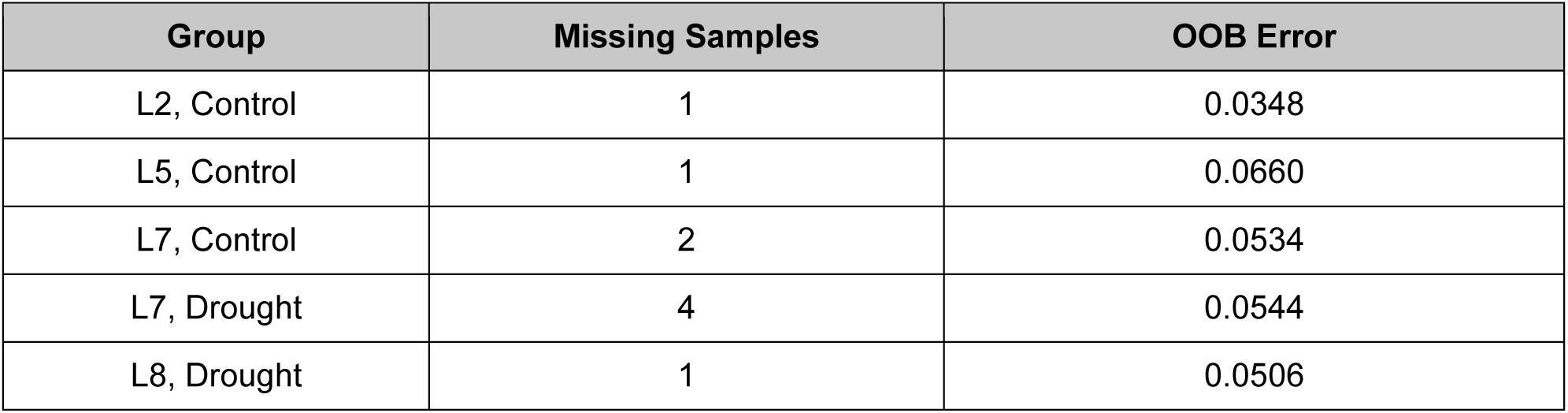
Out of bag (OOB) error of imputed missing values in harvest traits. In all samples, missing values were only in the total spike weight trait, except in L8, where total biomass was also missing. OOB error is calculated as normalized root mean square error (NRMSE) based on predictions for observed values that were left out of the training bootstrap samples (i.e., out-of-bag) when fitting the random forest.

| Group | Missing Samples | OOB Error |
| --- | --- | --- |
| L2, Control | 1 | 0.0348 |
| L5, Control | 1 | 0.0660 |
| L7, Control | 2 | 0.0534 |
| L7, Drought | 4 | 0.0544 |
| L8, Drought | 1 | 0.0506 |

**Supp. Table S4:** Outlier rate and re-imputation out of bag (OOB) error for each sensor type, including different protocols as described in Figure 1. Outliers were detected for each trait among all replicates within a genetic line within a treatment.

| Sensor type | Outlier Rate | Re-imputation OOB Error mean |
| --- | --- | --- |
| FC, Night protocol using conditional light | 0.0092 | 0.0194 ± 0.0322 |
| FC, Night protocol using high light | 0.0112 | 0.0209 ± 0.0337 |
| FC, Morning protocol for chlorophyll content | 0.0090 | 0.0563 ± 0.0867 |
| FC, Morning protocol for QY Lss | 0.0155 | 0.0330 ± 0.0403 |
| Thermal IR | 0.01 | 0.0166 ± 0.0114 |
| RGB | 0.0057 | 0.0168 ± 0.0485 |
| RGB color segmentation | 0.0147 | 0.1481 ± 0.2816 |
| HS, SWIR | 0.0073 | 0.0336 ± 0.0379 |
| HS, VNIR | 0.0075 | 0.0184 ± 0.0401 |
| Harvest | 0.0084 | 0.0315 ± 0.0499 |

**Supp. Table S5:** Rates of non-normality (NN) for each sensor type, including different protocols as described in Figure 1. Non-normality was tested for each trait within each data group, either separated by treatment or pooled across treatments. Groups identified as non-normal were corrected using a Box–Cox transformation, and NN rates were recalculated post-correction.

| Sensor type | NN rate separated | NN rate corrected separated | NN rate pooled | NN rate corrected pooled |
| --- | --- | --- | --- | --- |
| FC, Night protocol using conditional light | 0.0818 | 0.0389 | 0.0475 | 0.0278 |
| FC, Night protocol using high light | 0.0693 | 0.0302 | 0.0455 | 0.0236 |
| FC, Morning protocol for chlorophyll content | 0.1528 | 0.0250 | 0.0611 | 0.0167 |
| FC, Morning protocol for QY Lss | 0.1240 | 0.0600 | 0.0773 | 0.0693 |
| Thermal IR | 0.1370 | 0.1304 | 0.0348 | 0.0130 |
| RGB | 0.0434 | 0.0202 | 0.0694 | 0.0480 |
| RGB color segmentation | 0.0620 | 0.0480 | 0.0640 | 0.0520 |
| HS, SWIR | 0.0150 | 0.0100 | 0.0350 | 0.0100 |
| HS, VNIR | 0.0370 | 0.0290 | 0.0540 | 0.0380 |
| Harvest | 0.0577 | 0.0231 | 0.0846 | 0.0462 |

**Supp. Table S6.**
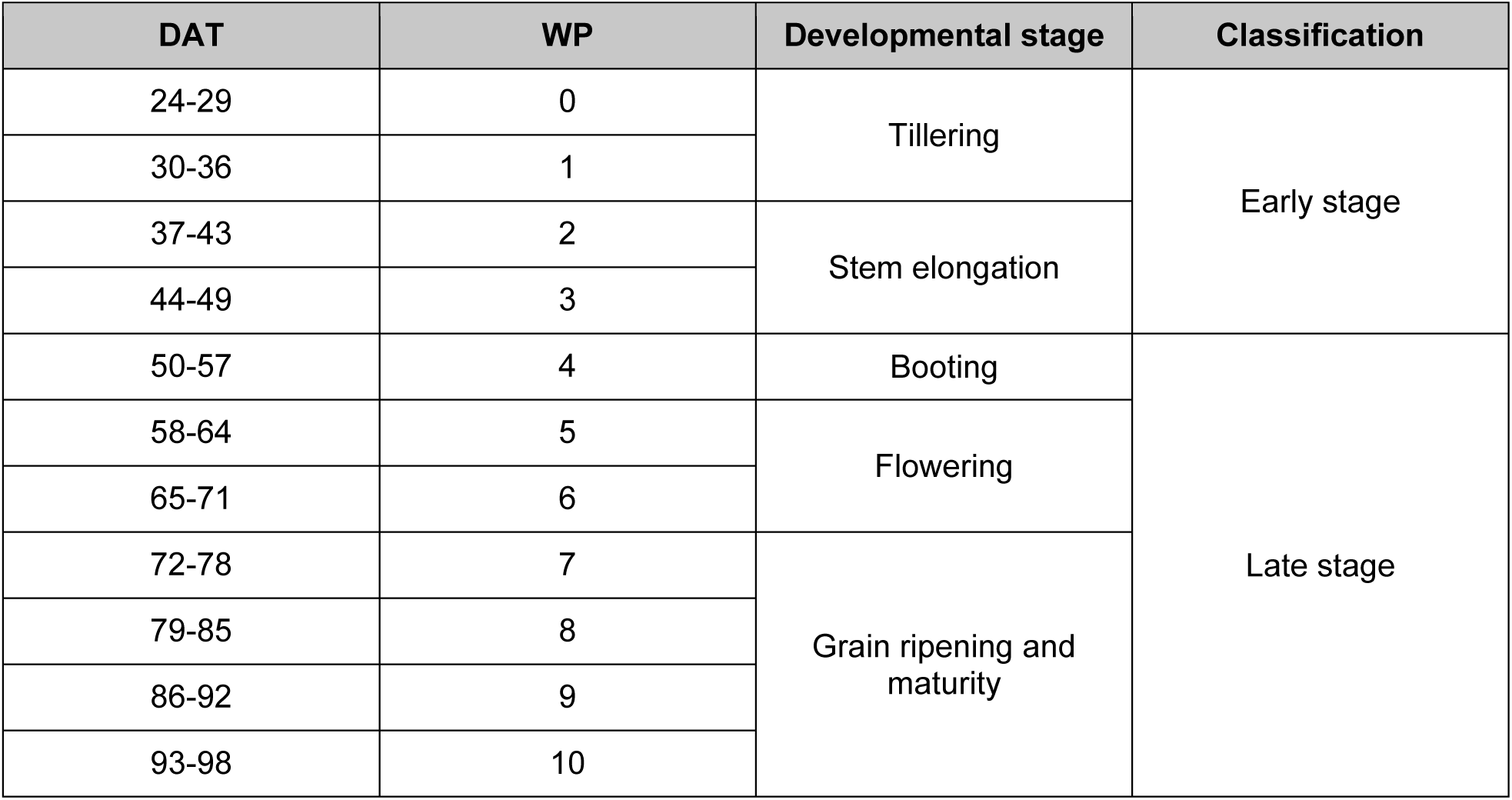
Mapping between the daily time point counting from days after transfer seedlings to light (DAT) for the non-aggregated dataset and the corresponding weekly phases (WP) for the aggregated dataset. The developmental stage was marked when most of the replicates among all the genetic lines reached a certain stage. Early stage refers to the vegetative and early reproductive phases, during which plants were exposed to 3 weeks of drought stress. Late stage refers to the late reproductive phase, when the flag leaf was fully expanded, and plants were exposed to a longer duration of drought stress.

| DAT | WP | Developmental stage | Classification |
| --- | --- | --- | --- |
| 24-29 | 0 | Tillering | Early stage |
| 30-36 | 1 |  |  |
| 37-43 | 2 | Stem elongation |  |
| 44-49 | 3 |  |  |
| 50-57 | 4 | Booting | Late stage |
| 58-64 | 5 | Flowering |  |
| 65-71 | 6 |  |  |
| 72-78 | 7 | Grain ripening and maturity |  |
| 79-85 | 8 |  |  |
| 86-92 | 9 |  |  |
| 93-98 | 10 |  |  |

**Supp. Table S7.** Performance of final LASSO models trained on the full predictor set using the optimal parameter (λ). Accuracies much higher than those found in the internal CV procedure during training suggest overfitting. “Training Group” refers to the data group used to train the model, while the R^2^ values show the accuracy when the model is applied to the data group in question.

| Response Trait | Training Group | R<sup>2</sup> Control | R<sup>2</sup> Drought | R<sup>2</sup> Pooled |
| --- | --- | --- | --- | --- |
| Biomass DW | Control | 0.935 | 0.876 | 0.894 |
| Biomass DW | Drought | 0.793 | 0.953 | 0.886 |
| Biomass DW | Pooled | 0.893 | 0.945 | 0.985 |
| Spike No | Control | 0.9 | 0.174 | 0.668 |
| Spike No | Drought | 0.272 | 0.822 | 0.539 |
| Spike No | Pooled | 0.852 | 0.821 | 0.898 |
| Total Spike W | Control | 0.958 | 0.483 | 0.849 |
| Total Spike W | Drought | 0.618 | 0.926 | 0.925 |
| Total Spike W | Pooled | 0.887 | 0.856 | 0.969 |
| Spike W.5Spk | Control | 0.785 | 0.649 | 0.654 |
| Spike W.5Spk | Drought | 0.589 | 0.816 | 0.587 |
| Spike W.5Spk | Pooled | 0.783 | 0.834 | 0.828 |

